# Retinoic acid coordinates the orderly construction of the mammalian body in the anterior-to-posterior sequence

**DOI:** 10.64898/2025.12.11.693671

**Authors:** Anita Banerjee, Sameera Krishna Yallapragada, Gabriel Torregrosa-Cortés, Bhakti J Vyas, Ramkumar Sambasivan

**Affiliations:** Indian Institute of Science Education and Research (IISER) Tirupati, Srinivasapuram, Panguru, Yerpedu mandal, Tirupati 517619, Andhra Pradesh, India; EMBL Barcelona, Dr. Aiguader, 88, PRBB Building, 08003 Barcelona, Spain; Institute for Stem Cell Science and Regenerative Medicine, GKVK-post, Bellary Road, Bengaluru 560065, Karnataka, India

## Abstract

In vertebrate embryos, anterior structures are formed early during gastrulation, while the posterior body develops subsequently. This temporal anterior-to-posterior developmental sequence is a fundamental aspect of animal development. Recent studies have shown that discrete regulatory modules involving Nodal and Wnt signaling pathways, in conjunction with T-box factors Eomes and Brachyury (Tbxt), underlie the differential specification of anterior and posterior mesoderm. However, the mechanism governing the transition from anterior to posterior mesoderm development remains unclear. Our findings suggest that retinoic acid signaling regulates the anterior-to-posterior developmental transition. Using mouse embryonic stem cell-based gastruloids, we show that the preclusion of retinoic acid signaling drives anterior mesoderm development, whereas the mesoderm specified in the presence of retinoic acid acquires posterior identity. Our observations indicate that retinoic acid signaling modulates the Wnt pathway. We demonstrate that both *Eomes* and *Tbxt* are essential for suppressing the posterior fate. The negative regulation of Wnt signaling by these T-box factors is critical, along with the preclusion of retinoic acid signaling, to ensure orderly anterior-posterior developmental progression. We propose that the preclusion of retinoic acid signaling early during gastrulation is crucial to maintain low Wnt signal levels, thereby driving anterior mesoderm specification. Subsequently, the activation of retinoic acid signaling results in high Wnt conditions, facilitating a switch to posterior mesoderm development.

## Introduction

The animal body is constructed in a head-to-tail sequence. During embryogenesis, the anterior structures are specified, followed by the progressive formation of the posterior body (Stern et al., 2006). The deep conservation observed across bilateria highlights the fundamental nature of this mode of development. Despite its fundamental nature, the mechanism of anterior-to-posterior (A-P) developmental progression remains poorly understood. Differentiation of the mesoderm germ layer along the anterior-posterior axis offers an excellent model for investigating A-P progression. Recently, there has been renewed efforts to elucidate the network regulating the differential specification of anterior and posterior mesoderm.

Nodal, a TGFβ superfamily member, and Wnt/β-catenin signaling determine the mesodermal fate of pluripotent epiblast cells in mammalian embryos. These signals induce the expression of T-box transcription factors Eomes and Tbxt (Brachyury or T), which regulate the mesodermal gene network (Ramkumar & Anderson, 2011). In mouse embyros, mesoderm emerging from the early primitive streak between embryonic days E6.5 and E7.5 takes anterior identity (Kinder et al., 1999; Lawson et al., 1991). After E7.5, the late primitive streak gives rise to posterior mesoderm. Eomes and Nodal signaling are essential for anterior mesoderm specification (Arnold et al., 2008; Costello et al., 2011; Simon et al., 2017; Van Den Ameele et al., 2012). Conversely, specification of the posterior mesoderm requires the function of Tbxt and Wnt3a (Herrmann, 1991; Martin & Kimelman, 2008a; Nandkishore et al., 2018; Smith et al., 1991; Takada et al., 1994; Yamaguchi, Takada, et al., 1999). Wnt/β-catenin signaling imparts posterior identity via the activation of caudal homeobox factors, specifically Cdx2 (Ikeya & Takada, 2001; Lickert et al., 2000; Pilon et al., 2006, 2007). The Wnt-Cdx axis appears to be a deeply conserved pathway governing embryonic posterior development across bilateria (Martin & Kimelman, 2009).

The transition from early to late streak is marked by a shift from *Wnt3* to *Wnt3a* expression (Yamaguchi, 2001), correlating with decreased *Eomes* and increased *Tbxt* expression in the newly formed mesoderm (Wehmeyer et al., 2025). Although *Eomes* and *Tbxt* are co-expressed in the early streak, Eomes restricts Tbxt function, promoting an anterior trajectory (Schüle et al., 2023). Recent studies have demonstrated that low Wnt signaling, established by the Nodal-Eomes-Wnt3 network, specify the anterior mesoderm, whereas high Wnt signaling, established by the Tbxt-Wnt3a network, specifies the posterior mesoderm (Dias et al., 2025; Wehmeyer et al., 2025). Furthermore, the activation of the positive feedback module of Wnt signaling and Cdx2, a caudal homeobox factor of the parahox cluster, signifies the transition to a posterior developmental programme (Amin et al., 2016; van de Ven et al., 2011; van Rooijen et al., 2012; Young et al., 2009; Young & Deschamps, 2009). These investigations led to a model of regulatory networks governing the differential specification of anterior and posterior mesoderm. However, the mechanism underlying the transition from the anterior to the posterior program remains elusive.

Retinoic acid (RA) is undetectable in early mouse gastrulae, and its synthesis begins around embryonic day 7.5 (E7.5) with the activation of Aldh1a2 in the differentiating paraxial mesoderm (Ang et al., 1996; Rossant et al., 1991). Notably, maternal RA is degraded in early streak embryos by Cyp26a enzymes to maintain a RA-negative environment (Uehara et al., 2009). Although the posterior mesoderm is specified, *Aldh1a2^-/-^* mouse embryos display severe reduction in the posterior body elongation (Niederreither et al., 1999). Moreover, it is well established that RA signaling regulates body axis patterning through the regulation of Hox genes (Nolte et al., 2019). While the correlation between the initiation of RA signaling and early-to-late streak transition is evident, the causality has not been investigated. Although the role of RA in axial patterning following the activation of the posterior program has been extensively studied, its potential function in specifying anterior identity or in the A-P developmental transition remains to be elucidated.

Using mouse embryonic stem cell-derived gastruloids and single-cell transcriptomics, we demonstrate that the inhibition of retinoic acid (RA) signaling permits the specification of anterior mesoderm, whereas the presence of RA leads to posterior mesoderm specification. In the absence of RA signaling, Wnt/β-catenin signaling is attenuated. Taken together with the recent findings that a high Wnt signaling threshold accelerates the transition to posterior mesoderm, our study indicates that RA regulates the A-P transition by modulating the Wnt signaling pathway. Furthermore, using genetics, we reveal that *Eomes* is essential for suppressing the Wnt-Cdx2 module to inhibit the posterior fate. Surprisingly, suppression of the posterior fate in the absence of RA also requires *Tbxt* function. In the absence of RA, increased Wnt signaling in *Tbxt* mutants promotes posterior neural fate. Since the activation of Wnt-Cdx2 in the absence of either *Eomes* or *Tbxt* overrides the need for RA signaling for posterior fate specification, we infer that RA functions upstream of the Wnt-Cdx module to regulate the A-P progression. Collectively, our findings suggest that the transition from an RA-negative early streak to an RA-active late-streak stage promotes the orderly A-P developmental transition.

## Results

### Preclusion of the RA signal specifies anterior mesoderm

Classical gastruloids are characterized by the presence of post-occipital embryonic tissues (Beccari et al., 2018; Turner et al., 2017). Although cardiomyocytes, which originate from the anterior mesoderm, have been successfully derived through the application of exogenous cardiogenic factors (Rossi et al., 2021), the mesoderm generated in these gastruloids predominantly consists of the posterior subtype, with limited specification of the anterior mesoderm (Mayran et al., 2025; Rekaik et al., 2023; Rosen et al., 2022; Suppinger et al., 2023; van den Brink et al., 2020). Hence, we refer to the classical gastruloids as posterior mesoderm (PM) gastruloids here. We modulated the signaling cues in this model to investigate the mechanism underlying the differential specification of anterior and posterior mesoderm.

Wnt/β-catenin, Nodal, and FGF signaling pathways are key regulators of early mesoderm development in the mouse gastrula (Arnold & Robertson, 2009; Ciruna & Rossant, 2001; Ramkumar & Anderson, 2011), and several studies have focused on these pathways. In contrast, the RA signaling pathway has been less extensively studied in this context. Studies have shown that RA is not synthesized in the embryos early on, and maternal RA is excluded during early mouse gastrulation when the anterior mesoderm emerges from the primitive streak (Ang et al., 1996; Rossant et al., 1991; Uehara et al., 2009). RA synthesis begins around embryonic day E7.5, coinciding with the transition to posterior mesoderm formation (Ang et al., 1996; Rossant et al., 1991). Analysis of the single-cell ATLAS of mouse gastrulation (Pijuan-Sala et al., 2019) revealed that *Aldh1a2* RNA, encoding a critical enzyme in RA biosynthesis from Vitamin A, could be barely detected in the primitive streak and nascent mesoderm of early gastrulae, whereas the transcripts of RA-degrading cytochrome oxidase enzyme *Cyp26A1* showed strong expression (Figure S1, A-B). Given the role of RA in posterior patterning (Durston et al., 1989; Kessel & Gruss, 1991; Ruiz I Altaba & Jessell, 1991), we hypothesized that exclusion of RA may be essential for specifying the anterior mesoderm. To test this hypothesis, we generated gastruloids in the absence of vitamin A, the precursor of RA (Figure 1A). Mouse embryonic stem cells (ESCs) were cultured in N2B27 medium, using a version of B27 devoid of vitamin A. Unlike PM gastruloids, these gastruloids showed limited elongation at 120 hours (h) (Figure 1, B-C). They exhibited robust expression of Isl1, a cardiopharyngeal mesoderm marker which is a subset of the anterior mesoderm (Figure 1B). Notably, the posterior pole, characterized by polarized Tbxt and Cdx2 expression, was absent. Tbxt was expressed in a nonpolar manner, and Cdx2 expression was greatly reduced and overlapped with that of Tbxt (Figure 1B, D). These observations suggest that inducing Wnt signaling in the absence of the RA signal specifies anterior mesoderm, and suppresses Cdx2 expression and posterior pole formation. Notably, the minus vitamin A gastruloids differentiated to generate beating cardiac organoids when the culture was continued beyond 144h in the presence of cardiogenic factors, FGF10 and IGF1 (Figure 1E). The efficient cardiac differentiation underscored the anterior identity of the mesoderm formed in these gastruloids. As these gastruloids form the anterior mesoderm, we designated them anterior mesoderm (AM) gastruloids.

**Figure 1.**
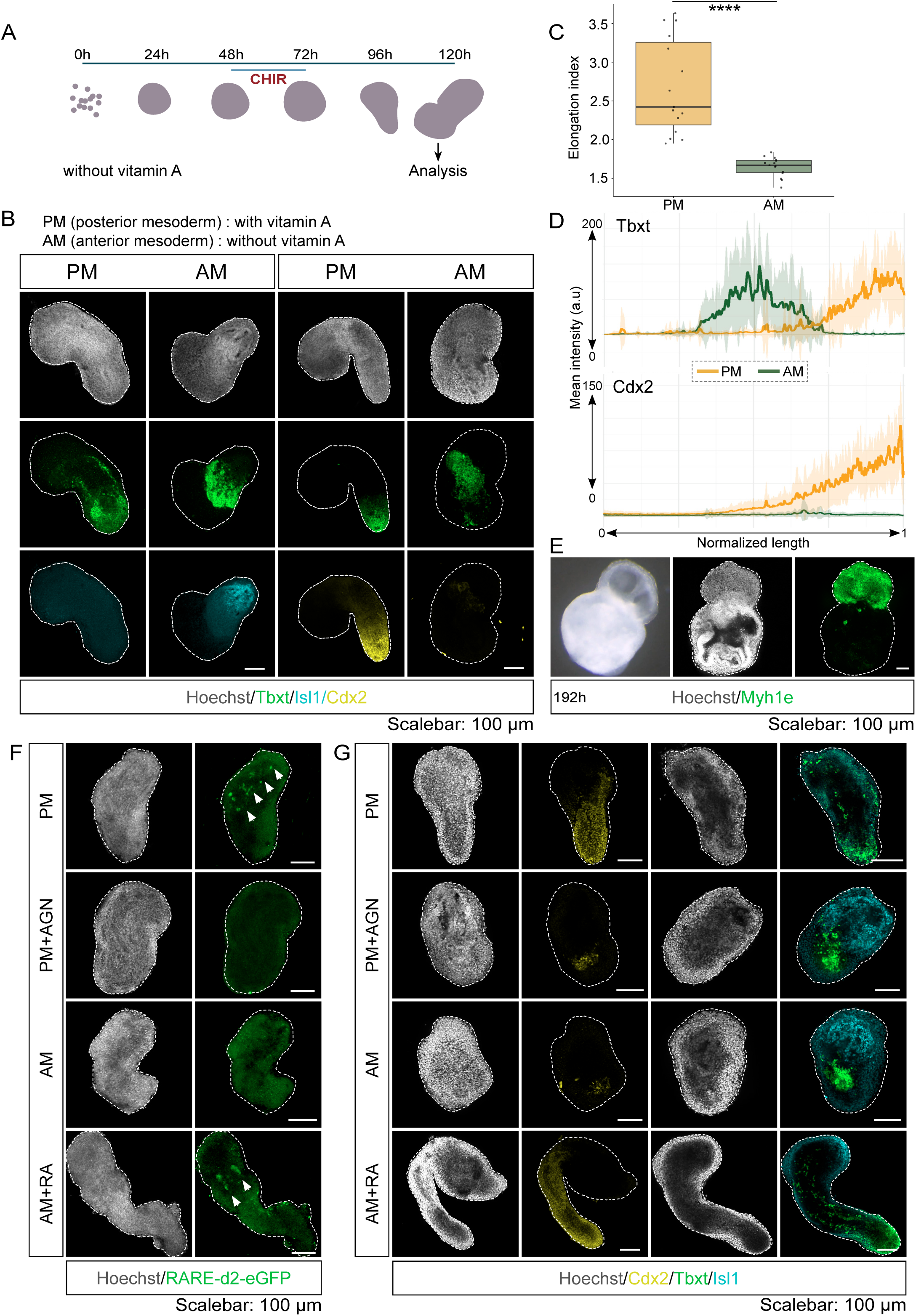
Removal of vitamin A or inhibition of retinoic acid signaling induces anterior mesoderm in gastruloids. (A) Schematic of the gastruloid generation protocol. To preclude retinoic acid (RA) signaling, B27 supplement without vitamin A, the biosynthetic precursor of RA, was used. CHIR - CHIR99021, a small molecule Wnt signal agonist. (B) Confocal images of gastruloids immunostained for the indicated markers. Tbxt (T/Brachyury); N = 6 experimental replicates, n = 50 PM gastruloids, and n = 66 AM gastruloids, posterior marker - Cdx2 (N = 3, n = 24 PM, and 24 AM gastruloids), anterior mesoderm marker - Isl1 (N = 3, n = 22 PM, and 28 AM gastruloids). (C) Comparison of cell elongation index between posterior mesoderm (PM) and anterior mesoderm (AM) gastruloids. Box plots show the median (central line), interquartile range (box), and individual data points. Paired t-test, n = 15 PM and 15 AM; ****p < 0.0001. (D) Line-scan quantification of Tbxt and Cdx2 expression along the long-axis of gastruloids from the experiment shown in (B). Intensity profiles along the length of gastruloids were obtained using the Fiji line segment tool, and gastruloid lengths were normalized between 0 and 1. The solid line traces the mean intensity (n=17 PM and 15 AM gastruloids), and the shaded region denotes the standard deviation. (E) Brightfield image and confocal images of immunostained RA-precluded gastruloids induced with a cardiogenic cocktail from 120 to 192 hours. Analysis at 192h revealed the formation of cardiac organoids expressing Myh14. Bright-field imaging reveals chambered organization in some of the cardiac organoids. (N = 4; n =15). (F) Confocal images of immunostained gastruloids. (N = 4, n = 23 PM, n = 25 PM + AGN, n = 23 AM, n = 26 AM + RA). AGN193109 (AGN) is an inverse agonist of retinoic acid receptors (RARs). Exogenous RA was added to AM gastruloids. (G) Confocal images of immunostained gastruloids. Isl1/Tbxt (N = 3, n = 20 AM, 14 AM + RA, 14 PM, 18 PM + AGN), Cdx2 (N = 3, n = 19 AM, 22 AM + RA, 23 PM, 17 PM + AGN). *Scale bar: 100 μm. All confocal images are represented as maximum intensity projections of z-stacks*.

Analysis of the RA signal reporter RARE-d2eGFP showed that RA signaling was undetectable in the minus vitamin A / AM gastruloids but was active in the posterior mesoderm (PM) gastruloids (Figure 1F). Furthermore, the inhibition of RA signaling through the use of an RA receptor antagonist AGN193109 during gastruloid generation replicated the AM-gastruloid phenotype (Figure 1, F-G). On the other hand, when RA is added exogenously, the ‘minus vitamin A’ gastruloids elongate and manifest polarized Tbxt and elevated Cdx2 expression similar to the standard ‘plus vitamin A’ PM gastruloids (Figure 1, F-G). These findings demonstrate that the RA signal is precluded when the culture environment lacks vitamin A. Collectively, these results underscore that the absence of the RA signal is essential for the specification of anterior mesoderm.

To investigate the mesodermal lineages present in AM gastruloids, we conducted single-cell transcriptome analysis. A total of 20,852 cells from 96 gastruloids each from three replicates were sequenced using the 10X Genomics platform. Through dimensional reduction and clustering analysis, we identified and annotated 15 distinct cell types within AM gastruloids (Figure 2A). Similar to PM gastruloids (Mayran et al., 2025; Rekaik et al., 2023; Rosen et al., 2022; Suppinger et al., 2023; van den Brink et al., 2020), mesodermal cell types predominate in AM gastruloids. However, in contrast to PM gastruloids, where the anterior mesoderm constitutes only a minor fraction (6.3%, as reported by (van den Brink et al., 2020)), anterior mesoderm-related clusters together with the prechordal plate cell type accounted for 61.6% of the cells in AM gastruloids (Figure 2B). The posterior mesoderm-related clusters, including somites, node/notochord, and caudal epiblast, comprised only 16.5% of the cells in the AM gastruloids. Based on established gene expression signatures, we identified the anterior mesoderm clusters as cardiopharyngeal mesoderm (*Isl1, Tbx1, Pitx2, Gata6,* and *Fgf10*), pharyngeal mesoderm 1 and 2 (*Isl1, Tbx1, Tcf21, Pitx2, Mesp1,* and *Lhx1*) (Figure 2C, S2A), and anterior paraxial mesoderm (*Tbx1, Meox1, Tcf15, Foxc2, Uncx, Hoxa2,* and *Hoxb2*) (Figure S2C). Notably, we identified one of the clusters as prechordal plate based on the expression of *Otx2, Hesx1, Six3, Sox17, Foxa2* and *Gsc* (Figure 2A, C, D, S2B). In addition to the prechordal plate, we identified a *Foxa2*+ and *Sox17*+ endoderm cluster, a subset of which exhibits pharyngeal identity (*Shh* (Moore-Scott & Manley, 2005)), *Gsc* (P. Thomas & Beddington, 1996), *Otx2* (Ee et al., 2025), *Eya1* and *Six1* (Zou et al., 2006), while a smaller subset is characterized by posterior identity (*Hoxb8* and *Cdx2*) (Figure S2D). The two clusters of differentiated somites, sclerotome and dermomyotome, lack cervical and thoracic Hox gene expression, leading us to consider them representative of differentiated occipital somites (see Figure S4A), which are part of the anterior mesoderm (Vyas et al., 2019; Wehmeyer et al., 2025). These findings clearly demonstrate that the preclusion of RA signaling specifies the anterior mesoderm.

**Figure 2.**
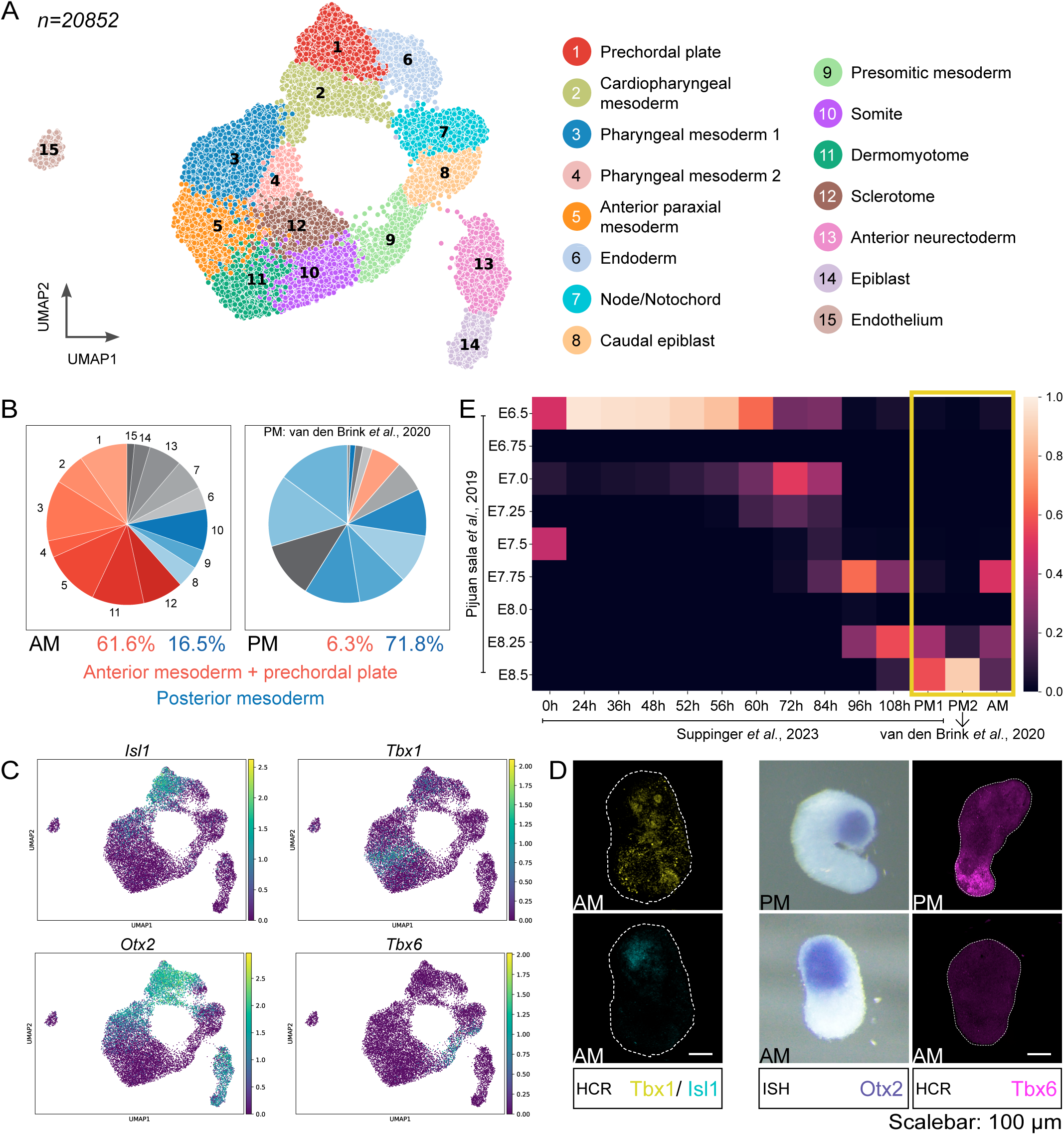
Single-cell transcriptomics reveal the anterior identity of RA signalling-precluded gastruloids. (A) UMAP plot of sc-RNA-seq data of 120h AM-gastruloids. *n=20852* cells obtained from 192 gastruloids pooled from 3 independent replicates. (B) Pie-chart comparing the relative proportions of anterior mesoderm (red) and posterior (blue) mesoderm-related clusters in AM gastruloids with those of PM gastruloids (van den Brink et al, 2020). Although it predominantly generates endoderm, we have included the prechordal plate cluster in the anterior mesoderm compartment, as it also gives rise to the prechordal mesoderm. (C) Gene interrogation plots for select marker genes: *Tbx1* and *Isl1* (cardiopharyngeal and pharyngeal mesoderm), *Otx2* (broad anterior), and *Tbx6* (posterior mesoderm). (D) Left panel: Confocal images of PM and AM gastruloids assayed using hybridization chain reaction (HCR). The broad domain of *Tbx1* expression and the spatially restricted *Isl1* expression (N = 3 experimental replicates, n = 20 gastruloids AM) are consistent with the single-cell sequencing data. Center panel: Brightfield images of gastruloids stained for *Otx2* mRNA by whole-mount in situ RNA hybridization (N = 3, n = 28 PM and 35 AM). Right panel: *Tbx6* HCR (N = 3, n = 11 PM and 11 AM). *Scale bar: 100 μm. All confocal images are represented as maximum intensity projections of z-stacks*. (E) Staging of AM gastruloids. An SCMAP projection of sc-seq data of AM gastruloids onto the mouse embryo single-cell gastrulation atlas to stage the gastruloids with respect to the embryonic age of mouse embryos. PM gastruloid datasets, 1 and 2, from the indicated references are also projected for comparison.

The ectoderm formed in AM gastruloids also exhibit anterior characteristics. The *Sox2*+ *Sox3*+ neuroectoderm cluster expresses *Hesx1* and *Otx2*, and lacks *Cdx2* and *Hox* gene expression, indicative of forebrain/midbrain identity (Martinez-Barbera et al., 2000; Simeone & Acampora, 2001; Sussel et al., 1999; P. Q. Thomas et al., 2001). It is likely that some of the epiblast cells are induced by the prechordal plate and pharyngeal endoderm to adopt the anterior neural fate. Consistent with the absence of a posterior pole, neuromesoderm progenitor-like population, and posterior neural cell types are absent in AM gastruloids. Therefore, our findings indicate that the preclusion of RA signaling predominantly specifies anterior identity across all germ layers.

Based on the expression profiles of *Cdx2*, *Nkx1-2*, *Tbxt*, and *Sox2*, in conjunction with a pluripotent signature, we have designated cluster #8 as caudal epiblast. However, given the low levels of Cdx2 protein and the non-polarized Tbxt expression (Figure 1D), we deduce that a posterior growth zone is not established in AM gastruloids (Gouti et al., 2014; Martin & Kimelman, 2008b; Neijts et al., 2014; Sambasivan & Steventon, 2021). This inference aligns with the observed low elongation index (Figure 1C) and the absence of spinal cord-like cells in the AM gastruloids. Moreover, *Tbx6*, a marker of the posterior paraxial mesoderm (Chapman & Papaioannou, 1998; Javali et al., 2017; Nowotschin et al., 2012; Takemoto et al., 2011) was undetectable by hybridization chain reaction (HCR) in AM gastruloids (Figure 2, C-D). Taken together, these observations suggest that the posterior program is suppressed in the absence of RA signaling.

### Preclusion of RA restrains the transition to posterior development

The transcriptome of PM gastruloids corresponds to that of E8.5 mouse embryos (Mayran et al., 2025; Suppinger et al., 2023; van den Brink et al., 2020), which fits with their posterior character. To stage AM gastruloids in relation to mouse embryos, we performed sc-map analysis to project the AM-gastruloid data onto the mouse gastrulation data (Pijuan-Sala et al., 2019). For comparative purposes, we included two PM-gastruloid datasets, one of which encompasses data across a time series (Suppinger et al., 2023; van den Brink et al., 2020). As previously reported, transcriptomically, the 120h PM gastruloids aligned with E8.5 embryos (Figure 2E). Notably, 120h AM gastruloids corresponded to an earlier embryonic stage (Figure 2E); they resembled E7.75 mouse embryos and 96h PM gastruloids. Time series analysis has shown that the trajectory of differentiation in gastruloids mirrors the embryonic temporal developmental sequence (Rosen et al., 2022; Suppinger et al., 2023). Consequently, AM gastruloids model the murine developmental window between E6.5 and E7.75. This observation suggests that the A-P developmental progression is either delayed or restrained in AM gastruloids than in PM gastruloids.

To investigate whether retinoic acid (RA) signaling influences the A-P progression, we conducted a comparative analysis of Hox gene expression in AM and PM gastruloids. The temporal collinearity of Hox gene expression functions as a developmental timer. Notably, sc-seq data revealed that the overall Hox gene expression levels were reduced and fewer cells expressed Hox genes in AM gastruloids (Figure 3A). Furthermore, cervical and thoracic Hox genes, such as *Hoxc5* to *Hoxc9*, are expressed in PM gastruloids but are scarcely expressed in AM gastruloids (Figure 3A). Although normalizing and comparing expression levels across datasets presents challenges, our RNA *in situ* hybridization (ISH) data corroborate the findings from the comparative sc-seq data analysis; classical ISH or hybridization chain reaction (HCR) for selected Hox genes confirmed the reduction in Hox gene expression both in terms of levels and the proportion of cells (Figure 3B). To further assess the A-P progression, we assayed the time course of anterior and posterior marker induction in AM versus PM gastruloids. Comparative analysis by RT-qPCR in the earlier time points of AM and PM gastruloids revealed elevated levels of anterior mesoderm genes *Eomes* and *Wnt3*, while the posterior mesoderm regulator *Wnt3a* remained at lower levels throughout in AM gastruloids (Figure 3C).

**Figure 3.**
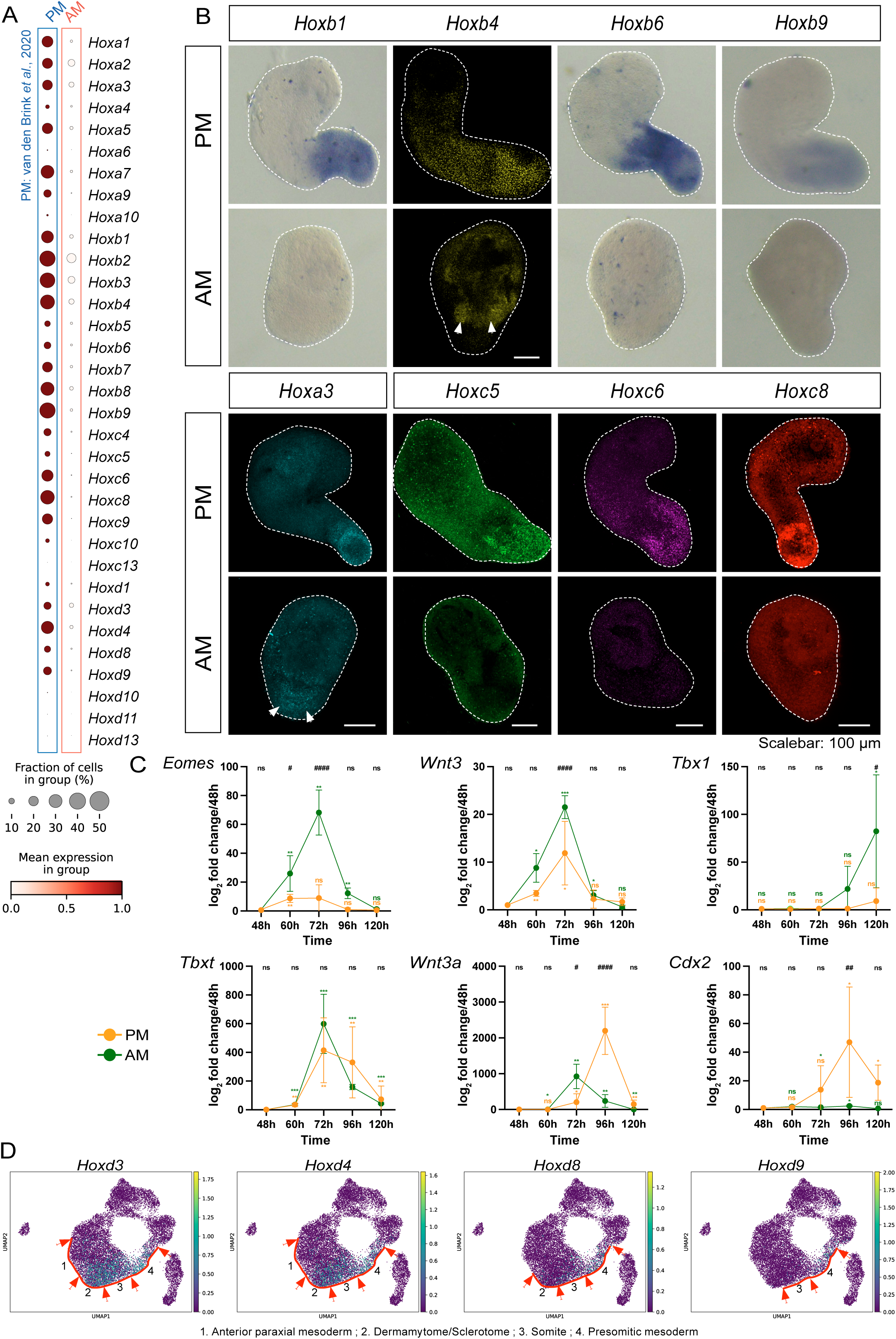
Transition to posterior program is repressed in AM gastruloids. (A) Dotplots of *Hox* gene expression in the merged PM (van den Brink et al., 2020) and AM single-cell datasets. Merging the datasets involves batch correction, which allows comparison. (B) Confocal and brightfield micrographs of gastruloids assayed by HCR and whole-mount chromogenic in situ hybridization (ISH), respectively. *Hoxb1* (N = 3 experimental replicates, n = 21 PM and 22 AM gastruloids), *Hoxb4* (N = 3, n = 10 PM and 9 AM), *Hoxb6* (N = 3, n = 21 PM and 25 AM), *Hoxb9* (N = 3, n = 25 PM and 19 AM), *Hoxa3* (N = 3, n = 15 PM and 16 AM), *Hoxc5* (N = 3, n = 17 PM and 17 AM), *Hoxc6* (N = 3, n = 11 PM and 11 AM), and *Hoxc8* (N = 3, n = 21 PM and 22 AM). *Scale bar: 100 μm. All confocal images are represented as maximum intensity projections of z-stacks*. (C) Comparison of gene expression trends across timepoints in PM and AM gastruloids. Data is represented as log₂ fold change relative to the 48h timepoint. Data represent mean ± SD of three biological replicates. Each replicate contained multiple gastruloids per time point (48h and 60h, n = 96 gastruloids; 72 and 96h, n = 72; 120h, n = 48). Post-hoc Sidak multiple comparisons, following a two-way ordinary ANOVA (see Methods), were performed between PM and AM at each time point. Statistical significance is indicated by hashtags (#) at the top of each graph. A two-tailed t-test was performed to compare changes over time within each condition. Significance is indicated by asterisks (*) above or below the error bars. Significance: *,# P ≤ 0.05; **,## P ≤ 0.01; ***,### P ≤ 0.001; ****,#### P ≤ 0.0001, ns = non-significant (P > 0.05). (D) Gene interrogation plots of the *Hoxd* genes showcase the spatial collinearity of *Hox* gene expression in AM gastruloids. *Hoxd3*, *Hoxd4* are enriched in differentiated somitic clusters, whereas *Hoxd8*, *Hoxd9* show restricted expression in undifferentiated somites and presomitic mesoderm. Red arrows indicate cluster boundaries. The red lines trace expression boundaries.

As previously reported for PM gastruloids (Beccari et al., 2018; Rekaik et al., 2023), spatial collinearity of Hox genes was also observed in AM gastruloids (Figure 3D). The spatial collinearity in AM gastruloids is depicted in the UMAP, within the A-P developmental gradient of the paraxial mesoderm; the differentiated somitic clusters of sclerotome and dermomyotome express 3’ Hox genes, while clusters representing undifferentiated somites, presomitic mesoderm, and caudal epiblast exhibit progressive expression of more 5’ Hox genes (Figure 3D). These observations support the anterior predominant identity of AM gastruloids and suggest that, despite the absence of a posterior pole and elongation, AM gastruloids are also organized along the A-P axis.

RA functions as an activator of Hox genes (Langston & Gudas, 1994; Nolte et al., 2019). Similarly, Cdx2 serves as a positive regulator of the middle Hox genes of the genomic cluster (Neijts et al., 2017; Rawat et al., 2008; van den Akker et al., 2002), which confers thoracic identity. Since Cdx2 expression is diminished when RA signaling is precluded, the dual block of Hox gene expression may serve as a mechanism for the orderly transition to a posterior developmental program. We propose that the RA signal operates as a temporal regulator; RA signaling expedites the transition to posterior mesoderm development, whereas its preclusion restrains the transition, thereby ensuring a temporal window for anterior mesoderm development.

### Wnt signaling is attenuated in the absence of RA signaling

The threshold of Wnt signaling paces the transition from anterior to posterior mesoderm development (Dias et al., 2025). Given that our findings suggest a similar role for RA signaling, we investigated whether RA signaling influences the Wnt pathway. To examine this, we compared Wnt signaling levels between AM and PM gastruloids using the *Tcf-Lef:mCherry* reporter (Faunes et al., 2013; Ferrer-Vaquer et al., 2010). Reporter expression was reduced in AM gastruloids by 96h (Figure 4A). Assaying by confocal microscopy and flow cytometry, we observed a significant decrease in the reporter signal intensity in AM gastruloids compared with that in PM gastruloids at 120h (Figure 4, A-B). This is striking because both cultures were exposed to the same pulse of Wnt agonist, 3μM CHIR99021, for 24h. The results with the reporter assay was corroborated by whole-mount hybridization chain reaction (HCR) for *Axin2*, a target of Wnt signaling (Jho et al., 2002), and *Dkk1*, which encodes a Wnt inhibitor (Niehrs et al., 2001). *Axin2* levels were comparable in AM and PM gastruloids at both 72h and 96h, but it was scarcely detectable in AM gastruloids at 120h (Figure 4C). Conversely, *Dkk1* was widely expressed at elevated levels within AM gastruloids. Notably, at 96 hours, *Axin2* and *Dkk1* RNA were localized at opposite poles (Figure 4C). Collectively, these results demonstrate that Wnt signaling is attenuated in the absence of RA signaling.

**Figure 4.**
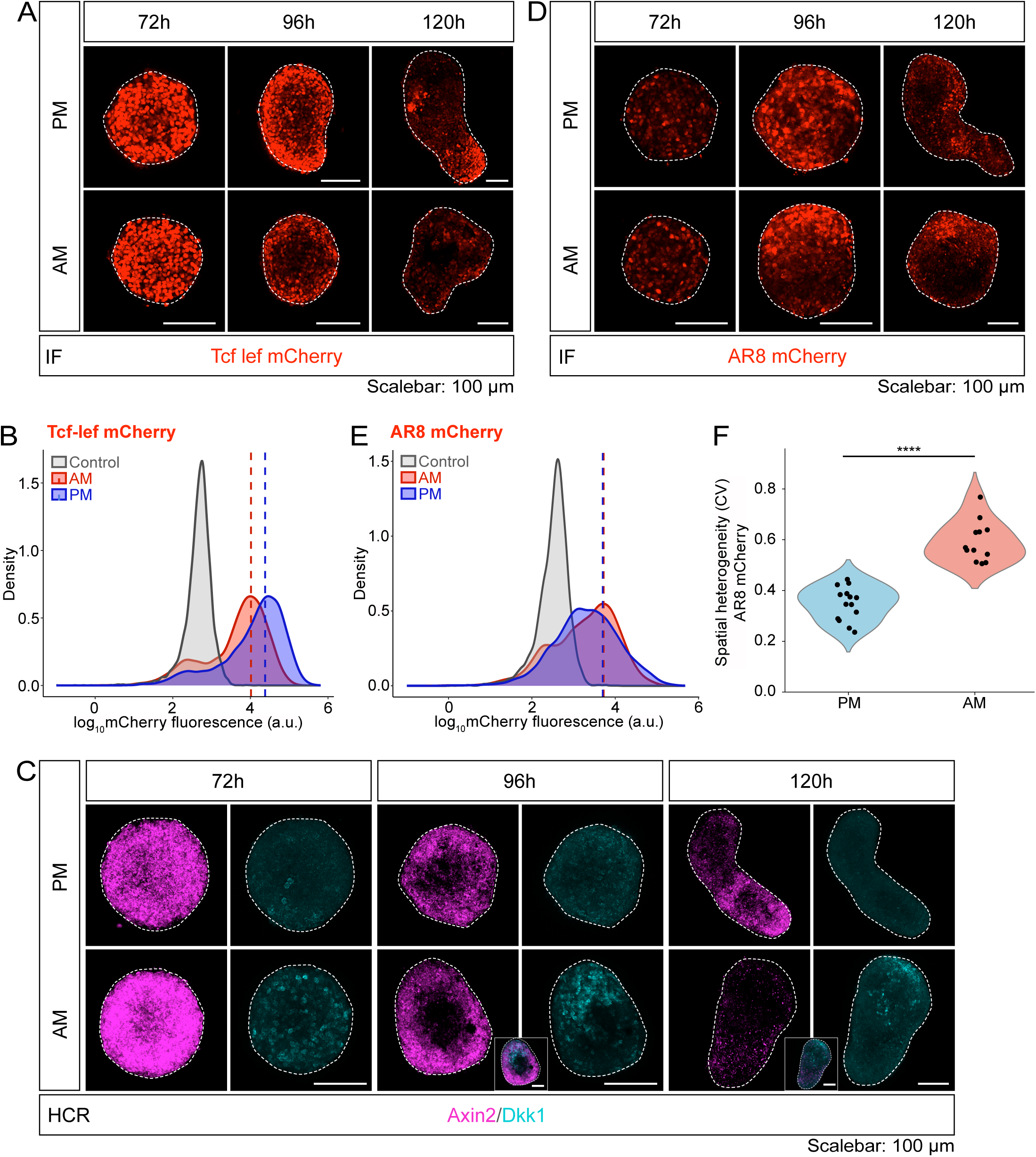
Absence of RA signaling attenuates Wnt signaling and restricts the Nodal signaling domain. (A) Confocal images of *Tcf-Lef:mCherry* reporter gastruloids immunostained with anti-mCherry antibody. (N = 3 experimental replicates, 72h (n = 20 PM and 19 AM gastruloids); 96h (n= 21 PM and 22 AM); 120h (n= 24 PM and 24 AM). (B) Flow cytometric analysis of 120h *Tcf-Lef:mCherry* reporter gastruloids. The positive threshold was set at 95% of cells from the negative control (unlabelled *E14Tg2a* mESCs). Dotted lines mark the median mCherry fluorescence of the samples. Similar results were obtained from 3 independent experiments; each sample is a pool of 48 individual gastruloids. (C) Confocal images of gastruloids assayed by HCR for *Axin2* (Wnt signalling target) and *Dkk1* (Wnt inhibitor). 72h (N = 3, n = 12 PM and 14 AM gastruloids); 120h (N = 4, n = 19 PM and 16 AM). Insets show the merge of both channels. (D) Confocal images of gastruloids immunostained for *AR8:mCherry* reporter expression 72h (N = 3, n = 17 PM, and 19 AM); 96h (N = 3, n = 19 PM, and 21 AM); 120h (N = 3, n = 27 PM, and 22 AM). *Scale bar: 100 μm. All confocal images are represented as maximum intensity projections of z-stacks*. (E) Flow analysis of 120h *AR8:mCherry* reporter gastruloids. Dotted lines indicate the median fluorescence intensity. (N = 3, 48 gastruloids pooled per replicate). (F) Quantification of spatial heterogeneity (coefficient of variation, CV) of *AR8:mCherry* expression in anterior mesoderm (AM) and posterior mesoderm (PM) gastruloids. Each point represents one gastruloid; violins indicate data distribution. Statistical significance was assessed using a two-sided Mann–Whitney U test (U = 168, p = 1.74 × 10⁻⁵).

Next, we investigated whether RA signaling modulated the Nodal pathway. Given that the induction of Nodal/Activin A signaling specifies the anterior mesoderm (Dias et al., 2025), we tested whether Nodal signaling is modulated in the absence of RA. Utilizing *AR8:mCherry*, a Nodal signaling reporter (McNamara et al., 2024; Serup et al., 2012), we observed that Nodal signaling levels were not affected in AM gastruloids compared to those in PM gastruloids (Figure 4, D-E). However, Nodal reporter expression was polarized in AM gastruloids, as reflected by a higher spatial heterogeneity, whereas PM gastruloids displayed a more uniform, dispersed expression pattern (Figure 4D, F). This contrasting spatial pattern of Nodal signaling is reminiscent of observations in mouse embryos. *Nodal* expression was restricted to the posterior epiblast at E6.25. Premature RA exposure disrupts this polarized Nodal expression, leading to ectopic expression in the anterior epiblast (Uehara et al., 2009). The expanded expression is driven by the intronic cis-regulatory element of the Nodal gene, which contains retinoic acid response element (RARE) sites. This observation provides additional evidence for the preclusion of RA signaling in the absence of vitamin A in the milieu.

The polarized Nodal signaling in the E6.25 mouse embryo is essential for protecting the anterior pole from posteriorization (Uehara et al., 2009). This is critical to create a Nodal-inhibited anterior pole, which promotes anterior mesoderm differentiation (Nandkishore et al., 2018). However, the significance of the spatial pattern of Nodal signaling for specifying anterior mesoderm within the primitive streak remains unknown.

### *Eomes* is necessary to suppress the posterior mesoderm fate

Wnt signaling activates the mesodermal program by targeting the T-box transcription factors, Eomes and Tbxt, which regulate the mesoderm gene network. To investigate the interaction between RA signaling and the Wnt-driven mesodermal network, we examined the function of *Eomes* and *Tbxt* in the absence of RA signaling. First, we studied the role of *Eomes*. Loss of *Eomes* function in mouse embryos results in failure of epithelial-to-mesenchymal transition in the primitive streak (Arnold et al., 2008; Russ et al., 2000), and *Eomes*-null mutant mESCs are unable to differentiate into cardiomyocytes, indicating a failure in specifying anterior mesodermal fate (Costello et al., 2011). We generated AM gastruloids (minus vitamin A gastruloids) using *Eomes^-/-^* mESCs and performed phenotypic analyses. Strikingly, unlike wild type AM gastruloids, *Eomes^-/-^* AM gastruloids expressed elevated levels of Cdx2; similarly to PM gastruloids, expression of Cdx2 and Tbxt was polarized (Figure 5, A-B). This is consistent with the recently reported observation that the *Eomes^-/-^*gastruloids generated through Activin A induction also initiate Cdx2 expression, whereas wild type Activin A gastruloids do not express Cdx2 (Wehmeyer et al., 2025). Although we observed variability across experiments, *Eomes^-/-^*AM gastruloids often displayed elongation. These observations indicate that the posterior growth zone is established even in the absence of RA signaling when *Eomes* function is lost.

**Figure 5.**
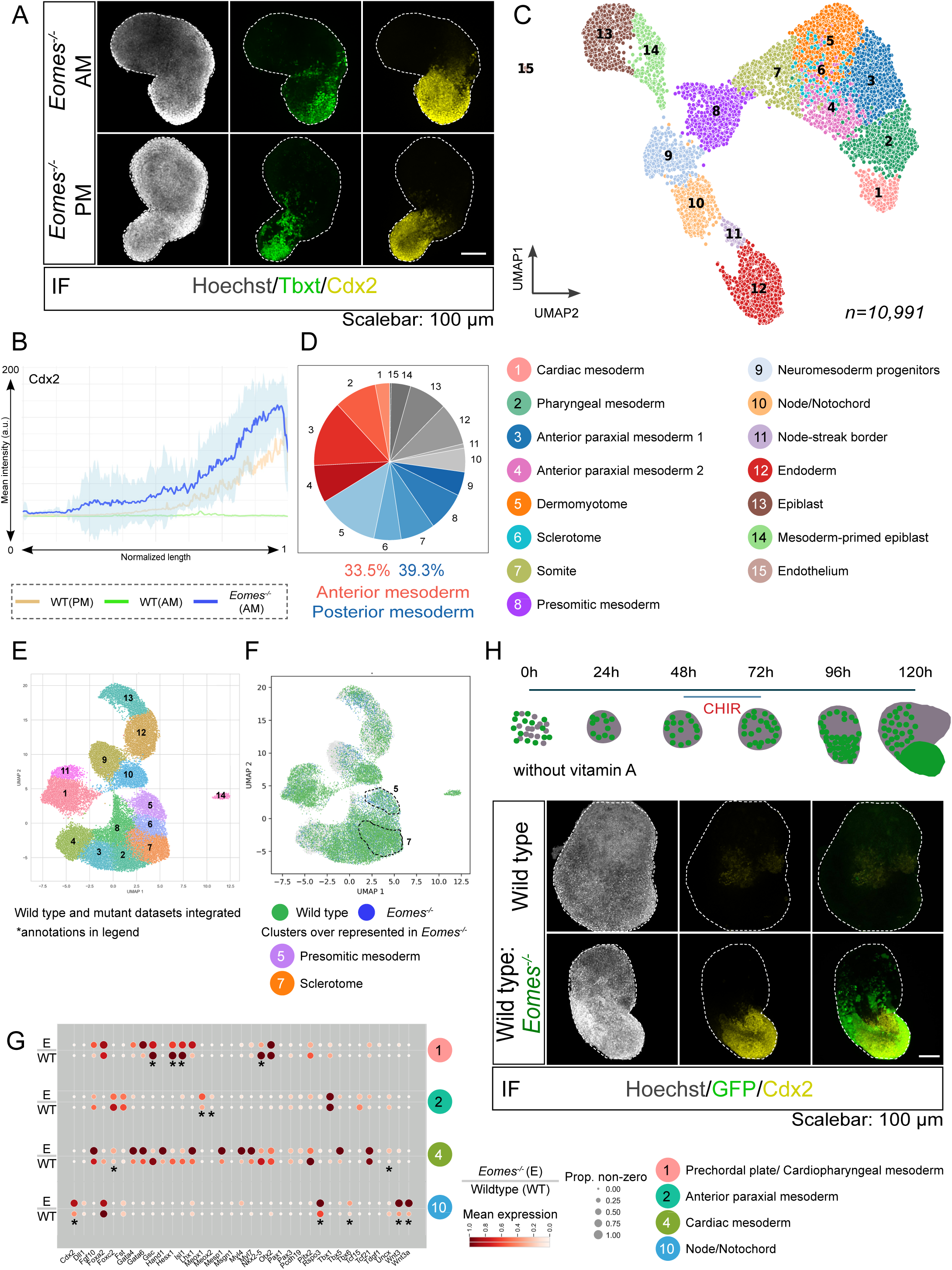
Posterior fate is induced in *Eomes* mutants despite the absence of RA signaling. (A) Confocal images of gastruloids immunostained for Tbxt (T/Brachyury) and posterior marker Cdx2. *Eomes^-/-^* (N = 5 experiments, n = 20 PM and 40 AM gastruloids). (B) Line-scan quantification of Cdx2 expression along the long-axis of *Eomes^-/-^* AM gastruloids (n = 16). Intensity profiles along the length of gastruloids were obtained using the Fiji line segment tool and gastruloid lengths of gastruloids were obtained using the Fiji line segment tool and gastruloids lengths were normalized between 0 and 1. The solid line indicates the mean intensity, and the shaded region denotes the standard deviation. The lighter orange and green mean intensity lines represent line-scan quantification of Cdx2 expression in wild type PM and AM gastruloids from Figure 1, shown here for comparison. (C) UMAP plot of sc-RNA-seq data of 120h *Eomes^-/-^* AM gastruloids. n = *10,991* cells obtained from 144 gastruloids pooled from 2 independent replicates. (D) Pie-chart comparing the relative proportions of anterior (red) and posterior (blue) mesoderm-related clusters in *Eomes^-^*^/-^ AM gastruloids. (E) UMAP plot of the integrated dataset of wild type, *Eomes*^-/-,^ and *Tbxt*^-/-^ AM gastruloids. The 14 clusters obtained were annotated based on the expression of marker genes. Annotations: 1-Prechordal plate & Cardiopharyngeal mesoderm, 2-Anterior paraxial mesoderm, 3-Pharyngeal mesoderm, 4-Cardiac mesoderm, 5-Presomitic mesoderm, 6-Somites, 7-Sclerotome, 8-Dermomyotome, 9-Neuromesoderm progenitors & Spinal cord, 10-Node/Notochord, 11-Endoderm, 12-Caudal epiblast, 13-Epiblast, 14-Endothelium. (F) UMAP showing co-embedding of wild type (green) and *Eomes^-/-^* (blue) AM gastruloid single-cell datasets. *Tbxt^-/-^* dataset is masked (grey). The clusters with a higher representation from the *Eomes^-/-^* dataset are highlighted with a dotted line. (G) The dot plot illustrates the expression of select marker genes in both wild type and *Eomes^-/-^*samples derived from the integrated single-cell RNA sequencing datasets. The color of the dots (red) signifies the average scaled gene expression, whereas the size of the dots indicates the percentage of cells expressing the gene within each cluster. Notable differences in expression are indicated by asterisks (*). Genes associated with the anterior regions, such as *Gsc, Hesx1, Isl1,* and *Nkx2-5*, exhibited reduced expression in *Eomes^-/-^* compared to the wild type, particularly within the prechordal plate/CPM cluster. Conversely, genes associated with the posterior regions showed increased expression in *Eomes^-/-^*, *Foxc2* and *Uncx,* in the cardiac mesoderm, *Meox1* and *Meox2* in the anterior paraxial mesoderm, and *Cdx2, Rspo3, Tbx6*, and *Wnt3a* in the node/notochord cluster. Confocal images of chimeric AM gastruloids, wild type:*Eomes*^-/-^-*GFP* in 50:50 ratio, immunostained for GFP and Cdx2. Wild type (N = 3, n = 31) and *Eomes^-/-^*-*GFP* (N = 3, n = 30). *Scale bar: 100 μm. All confocal images are represented as maximum intensity projections of z-stacks*.

To further investigate the phenotype, we conducted single-cell sequencing of *Eomes^-/-^*AM gastruloids. A total of 10,991 cells from 144 gastruloids pooled from two replicates were sequenced. UMAPs generated through dimension reduction and clustering analysis allowed the identification of 15 distinct cell types present in the mutant AM gastruloids (Figure 5C). Four anterior mesoderm clusters were identified: cardiac mesoderm, pharyngeal mesoderm, and anterior paraxial mesoderm 1 and 2 (Figure 5C, S3A). Notably, in comparison to the wild type, *Eomes* mutant AM gastruloids exhibited an increased proportion of posterior mesoderm cell types (Figure 5D, S3B). Unlike in the wild type, neuromesoderm progenitors, a major source of posterior paraxial mesoderm and spinal cord, were specified in the mutant gastruloids (Figure 5C, S3C). Furthermore, the posterior mesoderm clusters appeared to exhibit elevated levels of thoracic Hox gene expression relative to the wild type (Figure S3B, see also S4B). This posterior phenotype aligns with the increased, polarized Cdx2 expression and elongation observed in the mutants, and highlights the function of *Eomes* in inhibiting the posterior fate.

The sclerotome and dermomyotome clusters exhibit anterior/occipital identity in wild type AM gastruloids. In *Eomes^-/-^*AM gastruloids, these clusters show expression of thoracic *Hox* genes, such as *Hoxc5*, *Hoxc8* and *Hoxb8* (Figure S3B, S4B), and therefore they are of thoracic identity. We interpret the ‘posteriorized’ phenotype as an’anterior shift in the boundary’ of thoracic Hox genes into the cervical axial level. Despite the specification of the cardiac lineage and the detection of low levels of Isl1 RNA, Isl1 protein expression is reduced or undetectable in *Eomes*^-/-^ AM gastruloids (Figure S3A, D; (Costello et al., 2011)). These findings are consistent with the role of Eomes in restricting the function of Tbxt to curb posterior fate (Schüle et al., 2023).

To further address the differences in the cell types specified between wild type and *Eomes*^-/-^ AM gastruloids, we integrated the wild type and *Eomes^-/-^* datasets and generated a UMAP (Figure 5E). The clusters in the UMAP of the integrated dataset were annotated based on marker gene expression. Comparison of the distribution of *Eomes^-/-^*cells with those of wild type shows broadly similar composition of both types of AM gastruloids (Figure 5F). We observed an increased representation of presomitic mesoderm and sclerotome in the *Eomes^-/-^* (Figure 5F; see also Figure 6E). Moreover, the anterior mesoderm cell populations in the mutant AM gastruloids appear to be ‘posteriorized’. Analysis of differential expression of select genes showed diminished expression of anterior markers such as *Gsc, Hesx1*, and *Isl1*, and higher levels of posterior paraxial mesoderm markers, such as *Meox1, Meox2, Foxc2, Uncx* and *Tbx6* in the anterior mesoderm derivatives (Figure 5G). Notably, the posteriorization in the absence of *Eomes* function is evidenced by the preferential posterior localization of *Eomes^-/-^* cells in chimeric AM gastruloids generated by mixing GFP-labelled mutant and wild type cells (Figure 5H). Based on our findings, we conclude that the absence of RA signaling is permissive for anterior identity specification. However, anterior mesoderm specification is dependent on *Eomes* function. When RA signaling is precluded, *Eomes* functions to prevent the precocious posterior program activation to ensure orderly A-P developmental progression.

**Figure 6.**
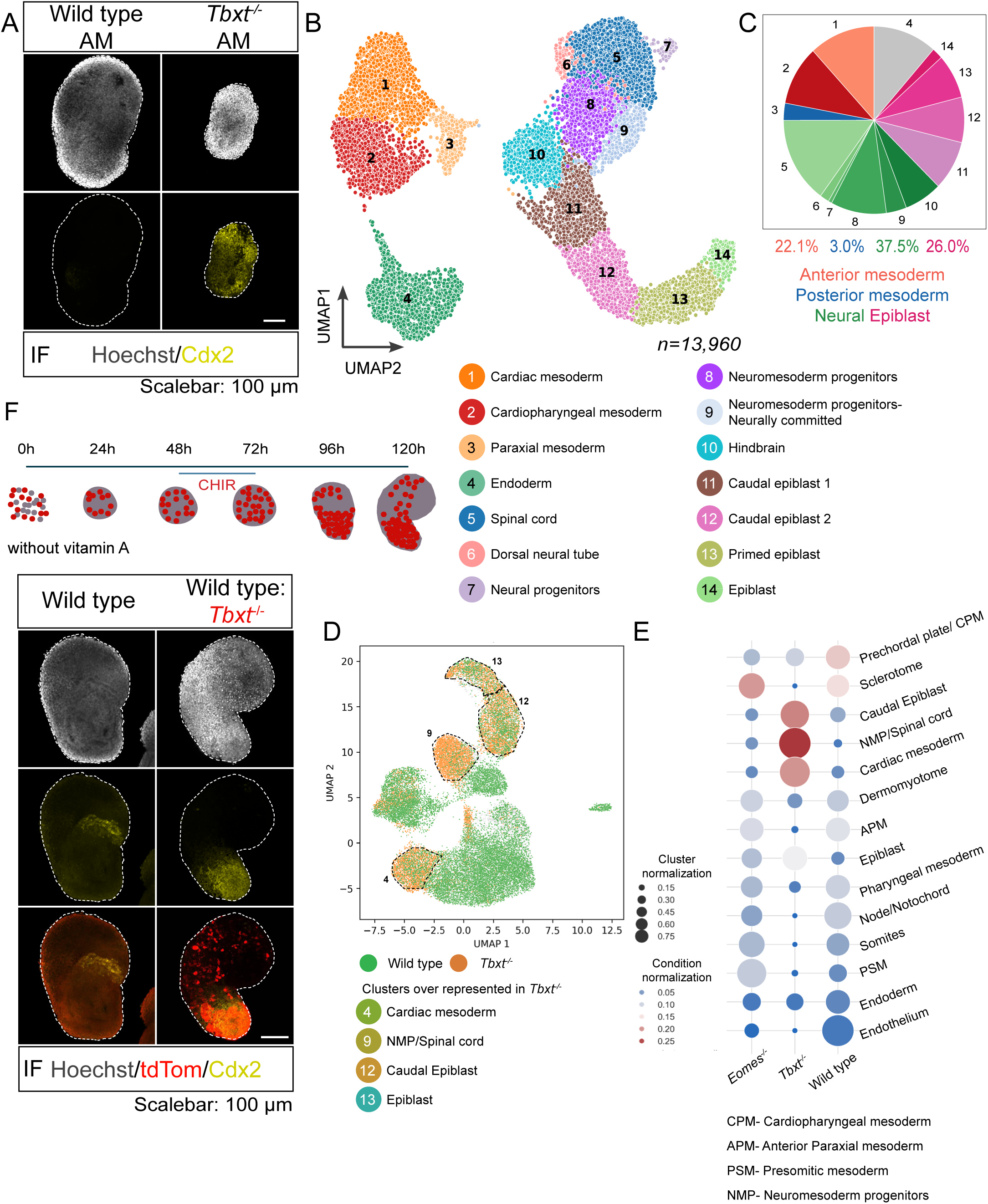
Loss of function of *Tbxt* activates posterior neural fate despite the absence of RA signaling. (A) Confocal images of gastruloids immunostained for the posterior marker Cdx2. Wild type (N = 3 experimental replicates, n = 16 AM gastruloids), *Tbxt^-/-^* (N = 3, n = 15 AM gastruloids). (B) UMAP plot of sc-RNA-seq data of 120h gastruloids *Tbxt^-/-^* AM gastruloids. *n=13,960* cells obtained from 192 gastruloids pooled from 2 independent replicates. (C) Pie-chart comparing the relative proportions of anterior (red) and posterior (blue) mesoderm-related, neural (green) and epiblast (pink) clusters in *Tbxt^-/-^* AM gastruloids. (D) UMAP showing co-embedding of wild type (green) and *Tbxt^-/-^* (orange) AM gastruloid datasets. *Eomes^-/-^* dataset is masked (grey). The clusters with a higher representation from the *Tbxt^-/-^* dataset are highlighted with a dotted line. (E) A clustermap comparing the proportion of cells in different clusters within (dot size) and across (dot colour) wild type, *Eomes^-/-^*and *Tbxt^-/-^* datasets. It was generated after the integration of the datasets. (F) Confocal images of chimeric AM gastruloids, wild type:*Tbxt*^-/-^-*tdTomato* in 50:50 ratio, immunostained for tdTomato and Cdx2. wild type (N = 3, n = 13) and *Tbxt^-/-^-tdTomato* (N = 3, n = 28). *Scale bar: 100 μm. All confocal images are represented as maximum intensity projections of z-stacks*.

### *Tbxt* suppresses posterior fate in the absence of RA

In contrast to *Eomes* mutants, anterior mesoderm specification and differentiation remain unaffected in *Tbxt* mutant mouse embryos; however, posterior mesoderm development is disrupted (Herrmann, 1991; Nandkishore et al., 2018). While *Tbxt* is expressed in the anterior mesoderm progenitors, its function in their specification is unclear. We investigated the function of *Tbxt* in anterior mesoderm development using AM gastruloid model. Unexpectedly, we observed elevated Cdx2 expression in *Τbxt^-/-^*AM gastruloids (Figure 6A) compared to the wild type. Although Cdx2 levels are increased, *Τbxt^-/-^* AM gastruloids do not elongate; they retain a spherical morphology. The activation of the posterior factor Cdx2 in *Τbxt^-/-^* gastruloids, despite the absence of RA signaling, presents a paradox given the critical role of *Tbxt* in the development of posterior mesoderm.

To investigate this unexpected mutant phenotype and the role of *Tbxt* in anterior developmental program, we conducted single-cell sequencing on *Τbxt^-/-^* AM gastruloids. A total of 13,960 cells from two experimental replicates of 96 gastruloids each were sequenced. UMAP visualization enabled the identification of 13 distinct cell types within *Τbxt^-/-^* AM gastruloids (Figure 6B). Consistent with existing literature, posterior paraxial mesoderm and its derivatives were nearly absent in 120h *Τbxt^-/-^*AM gastruloids. As expected, the anterior mesoderm cell types were specified; however, unlike the wild type AM and *Eomes* mutant AM gastruloids, *Τbxt^-/-^* AM gastruloids were predominantly composed of neural cells (46.1%) (Figure 6, B-C), with a reduced proportion of mesoderm (25%). The integration of our wild type, *Eomes^-/-^* and *Τbxt^-/-^* sc-seq datasets showed that *Τbxt^-/-^* AM gastruloids are relatively enriched for early cardiac mesoderm (Figure 6, D-E). This analysis also highlights the increased presence of posterior neural cells and epiblast-like cells uniquely in *Τbxt^-/-^*(Figure 6, D-E). These observations highlight the key role of *Tbxt* in mesoderm development.

Another striking phenotype of *Τbxt^-/-^* AM gastruloids is that the neural lineages were of posterior identity, unlike those in the wild type. The largest cluster in the UMAP was composed of spinal cord cells expressing *Pax6, Sox2,* and *Nkx1-2* (Figure 6B, S5). Two of the clusters corresponded to the neuromesoderm progenitors. Moreover, instead of the forebrain/midbrain neural cells specified in the wild type, *Τbxt^-/-^* AM gastruloids contained a hindbrain-like cluster expressing *Krox20* and *Phox2* (Figure 6B, S5). Hox gene expression further corroborated the posterior character of the mutant neuroectoderm (Figure S4C, S5). Therefore, a notable phenotype of *Τbxt^-/-^* AM is the induction of posterior fates, specifically the neuromesoderm progenitors and spinal cord, even in the absence of the RA signal. The expression of Cdx2 protein observed in *Τbxt^-/-^* gastruloids appears predominantly to be from the posterior neural cell types, spinal cord and neuromesoderm (Figure S5). Therefore, our findings show that the loss of *Tbxt* in conjunction with the absence of RA signalling drives posterior fate. Indeed, when we combined wild type and tdTomato-labelled *Τbxt^-/-^* mESCs, the resulting chimeric AM gastruloids exhibited elongation and polarized Cdx2 expression. In this context, the mutant cells predominantly localized at the posterior pole (Figure 6F), indicating that the conditions resulting from the loss of *Tbxt* and the absence of RA facilitated the development of posterior identity. Therefore, our findings demonstrate that Cdx2 expression and posterior fate are activated when *Tbxt* and the RA signal are absent.

### Wnt-Cdx2 posterior developmental module is suppressed in the absence of RA

When projected on to the single-cell transcriptomic data of mouse gastrula, the transcriptome of both *Eomes^-/-^* and *Τbxt^-/-^*AM gastruloids mapped partially to E8.5 embryo, further supporting the specification of posterior fates in the mutants (Figure 7A). To address the mechanism underlying the elevated expression of Cdx2 and the specification of posterior fate in T-box mutant gastruloids, despite the absence of retinoic acid (RA) signaling, we evaluated the levels of Wnt signaling in these mutant gastruloids. Canonical Wnt signaling is a principal regulator of Cdx2 (Chawengsaksophak et al., 2004; Wang & Shashikant, 2007) within the posterior axial progenitor pool. *Wnt3* is activated, but *Wnt3a* is suppressed by Eomes (Wehmeyer et al., 2025). In contrast, Tbxt and Wnt3a positively regulate each other through feedback mechanisms (Martin & Kimelman, 2008a; Wehmeyer et al., 2025; Yamaguchi, Yoshikawa, et al., 1999). First, we assessed the expression of *Wnt3* and *Wnt3a* in our sc-seq datasets (Figure 7B). In both the wild type and *Eomes^-/-^*, *Wnt3* and *Wnt3a* expression is confined primarily to the caudal epiblast, node/notochord or neuromesoderm cell clusters. Interestingly, the expression of *Wnt3* appeared derepressed in *Τbxt^-/-^* gastruloids, and it was widely expressed in the anterior mesoderm populations, in addition to the caudal epiblast-like population. As expected, *Wnt3a* expression was diminished in *Τbxt^-/-^* gastruloids (Figure 7B). The positive feedback loop of *Tbxt* and *Wnt3a* explains the *Wnt3a* downregulation in the mutants. This observation suggests that *Tbxt* suppresses *Wnt3* expression in the anterior mesoderm lineage.

**Figure 7.**
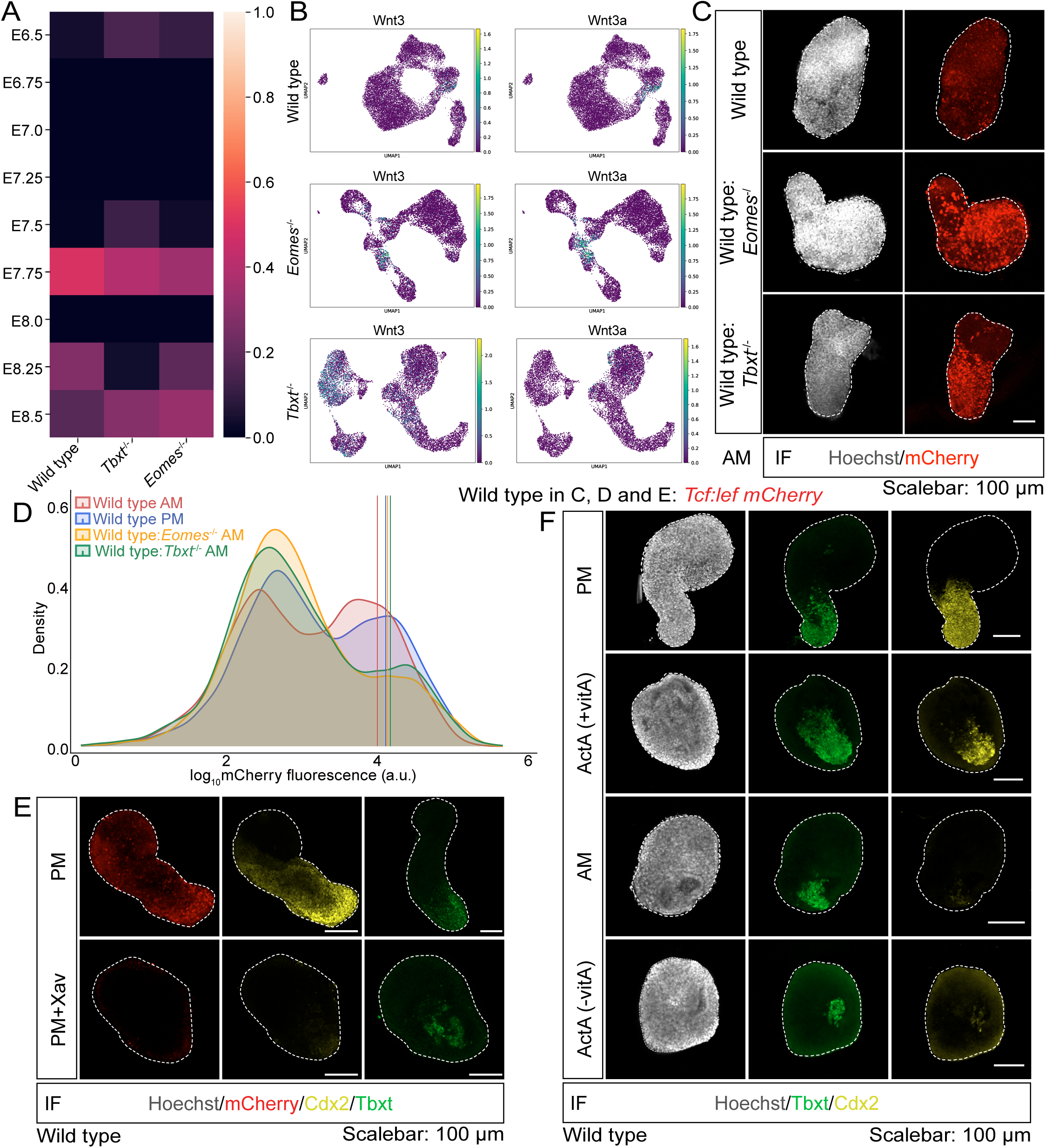
***Eomes* and *Tbxt* suppress the Wnt-Cdx2 module in the context of anterior mesoderm specification** (A) SCMAP projection of *Eomes^-/-^*, *Tbxt^-/-,^* and wild type AM gastruloid single cell sequencing data onto the mouse gastrulation atlas to assess the developmental age of the mutant gastruloids. (B) Gene interrogation plots of *Wnt3* and *Wnt3a* in wild type, *Eomes^-/-^*, and *Tbxt^-/-^* single-cell UMAPs. (C) Confocal images showing mCherry expression in wild type, *Eomes^-/-^,* and *Tbxt^-/-^* chimeric AM gastruloids. *Tcf-lef:mCherry* reporter was used as the wild type. Chimeric gastruloids generated by mixing wild type and mutant cells in a 50:50 ratio during plating (N = 3 experimental replicates, n = 21 wild type; 40 wild type:*Eomes^-/-^*; 27 wild type:*Tbxt^-/-^* gastruloids). (D) Flow-cytometric quantification of Wnt reporter activity in wild type (*Tcf-lef:mCherry* reporter) and chimeric AM gastruloids. Median mCherry fluorescence intensity of the *Tcf-Lef:mCherry* reporter was measured in wild type gastruloids under AM and PM conditions, as well as in chimeric AM gastruloids containing wild type and *Eomes^-/-^* or *Tbxt^-/-^* cells in a 50:50 ratio. Median intensities were as follows: wild type (AM): 3.983640; wild type (PM): 4.094799; wild type:Eomes^-/-^ (AM): 4.119460; wild type:*Tbxt*^-/-^ (AM): 4.157401. (E) Confocal microscopy images of indicated markers in PM and PM induced with Xav939 (Wnt inhibitor) from 72 to 120h in *Tcf-Lef:mCherry* reporter gastruloids. mCherry/Cdx2 (N = 3, n = 17 PM and 14 PM+Xav), Tbxt (N = 3, n = 19 PM and 14 PM+Xav). (F) Confocal imaging of indicated markers upon Activin-A (ActA) substitution for CHIR. (N = 5; n = 31 PM, 37 AM, 27 PM (+ActA), and 37 AM (+ActA)). *Scale bar: 100 μm. All confocal images are represented as maximum intensity projections of z-stacks*.

To assay Wnt signaling in the mutant gastruloids, we generated chimeric AM gastruloids by seeding a 50:50 mixture of *Tcf-Lef:mCherry* and *Eomes^-/-^* or *Tcf-Lef:mCherry* and *Τbxt^-/-^* mESCs. If the Wnt ligands are upregulated in the mutant cells, the wild type reporter cells within the chimera will provide the signaling read-out. Through confocal imaging and flow cytometry, we observed an increase in Wnt reporter expression in both *Eomes^-/-^* and *Τbxt^-/-^*chimeric gastruloids compared to non-chimeric wild type (*Tcf-Lef:mCherry*) AM gastruloids (Figure 7, C-D). These findings support the possibility that the enhanced Cdx2 expression in the mutants is due to the elevated Wnt signaling.

To determine whether the activation of Cdx2 can be attributed to increased Wnt signaling, we investigated the dependency of Cdx2 activation on Wnt signaling. PM gastruloids were treated with Xav939, a tankyrase inhibitor that suppresses the Wnt pathway, and Cdx2 levels were assessed by immunostaining. We observed that inhibition of Wnt signaling downregulated the expression of Cdx2 in wild type PM gastruloids (Figure 7E). Moreover, Xav939-treated PM gastruloids showed limited elongation and non-polar Tbxt expression akin to wild type AM gastruloids. These observations support the hypothesis that elevated Wnt signaling induces the activation of the Wnt-Cdx2 module, and that the de-repression of the Wnt-Cdx2 module underlies the accelerated acquisition of posterior fate in T-box factor mutant gastruloids, even in the absence of RA signaling.

To further probe the role of RA signaling in regulating Cdx2, we tested the expression of Cdx2 in the Activin A-based gastruloids. High concentrations of Activin A (ActA), an activator of the Nodal signaling pathway, drives anterior mesoderm specification (Dias et al., 2025). We assayed Cdx2 expression in high ActA gastruloids in presence/absence of vitamin A. Cdx2 is expressed in fewer cells in ActA gastruloids generated in the presence of vitamin A compared to PM gastruloids. However, without vitamin A, ActA gastruloids display a further reduction in Cdx2 levels (Figure 7F). This suggests that the preclusion of RA signaling is critical for effective suppression of Wnt-Cdx module and thus, the posterior fate.

## Discussion

The mechanisms underlying the sequential development of the animal body plan from the anterior to the posterior are beginning to be elucidated. Our research indicates that RA signaling plays a crucial role in facilitating the A-P developmental transition. Specifically, we propose that the absence of RA signaling during early gastrulation attenuates Wnt signaling, permitting the specification of anterior identity, including the anterior mesoderm. The Wnt-Cdx module is not activated at low Wnt signal thresholds, preventing premature activation of the posterior program. The subsequent activation of RA signaling during later gastrulation promotes increased Wnt signaling, which stabilizes the Wnt-Cdx module, causing a shift from the anterior to the posterior developmental program. Our findings indicate that RA signaling is integral to the regulatory logic governing the orderly A-P progression. The model emerging from recent research posits that the Nodal-Eomes-Wnt3 axis facilitates anterior mesoderm development, whereas the opposing Wnt3a-Tbxt axis promotes posterior mesoderm fate (Dias et al., 2025; Wehmeyer et al., 2025). Based on the evidence in the literature and our findings, we propose that RA signaling regulates the developmental transition from anterior to posterior mesoderm by impinging on the core anterior (Nodal-Eomes) and posterior (Wnt-Tbxt) networks (Figure 8).

**Figure 8.**
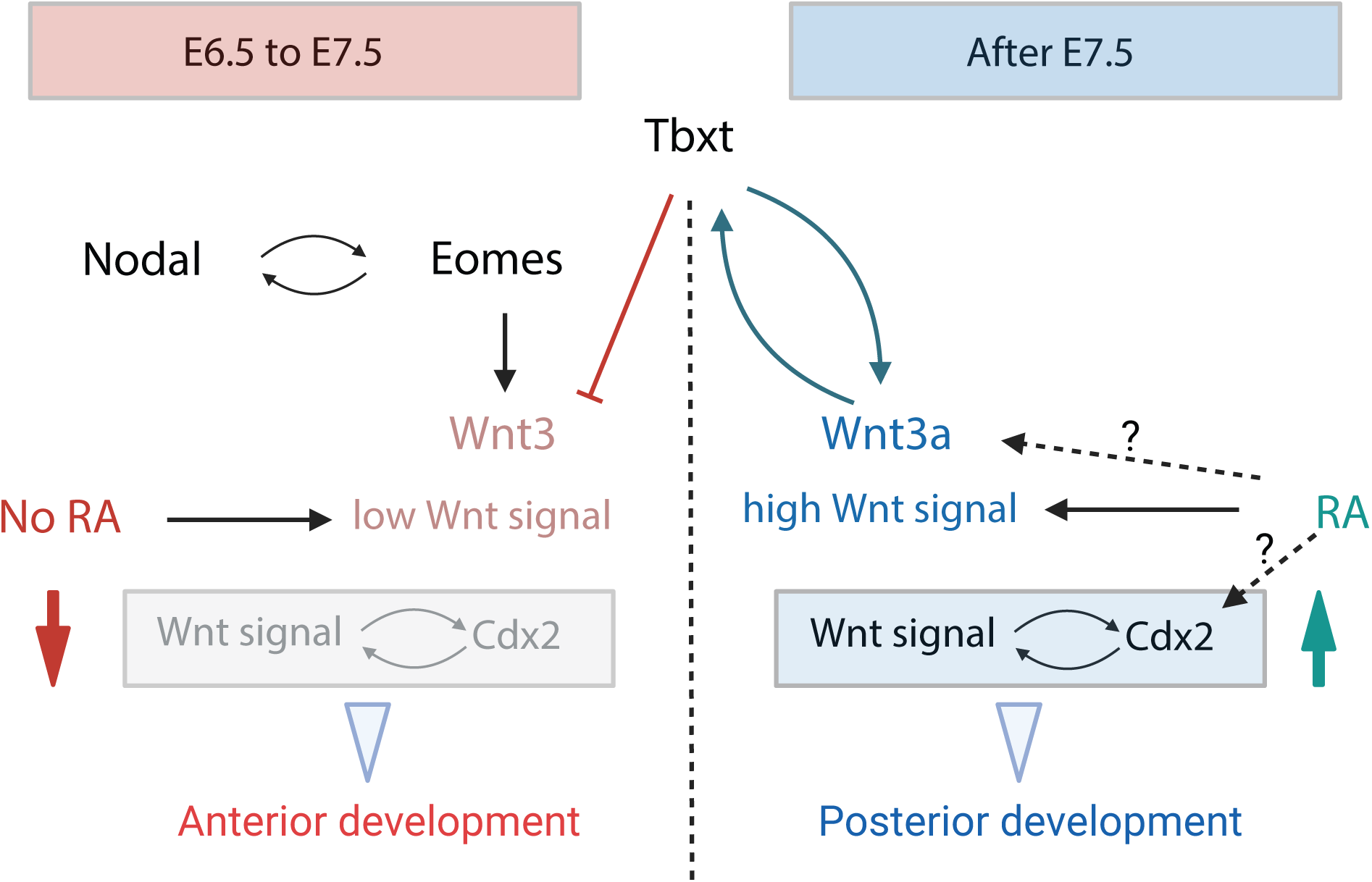
RA signalling regulates the orderly anterior to posterior transition of the mesoderm developmental program. Model summarizing the role of RA signaling in the mesoderm gene regulatory network governing the orderly developmental progression from anterior to posterior program. Between E6.5–E7.5, in the absence of RA, activation of the posterior program is prevented by maintaining Wnt signalling at a lower level. During this period, *Eomes* and *Tbxt* independently restrict premature posterior specification by downregulating Wnt activity through Wnt3a and Wnt3, respectively, thereby preventing activation of the Wnt–Cdx2 module and supporting anterior development. After E7.5, when RA signalling becomes active, a higher threshold of Wnt signalling is achieved by stabilizing the Tbxt–Wnt3a–Cdx2 module, which promotes posterior specification. Dashed arrows denote predicted RA-dependent regulation of Wnt3a and Cdx2.

*Eomes* and *Tbxt* are expressed in early streak mouse embryos, where they attenuate Wnt signaling by repressing *Wnt3a* and *Wnt3* expression, respectively (Wehmeyer et al., 2025). During the late streak stage, *Eomes* levels decrease, relieving its restriction on *Tbxt* function (Schüle et al., 2023), leading to elevated *Wnt3a* expression through positive feedback with *Tbxt* (Evans et al., 2012; Martin & Kimelman, 2008a; Schwaiger et al., 2022; Turner et al., 2014; Yamaguchi, Takada, et al., 1999). This may lead to low Wnt signaling in early streak embryos and heightened Wnt signaling in late streak embryos. Since Cdx2 is a Wnt signal target (Amin et al., 2016), we speculate that high Wnt signal threshold induces Cdx2 expression, activating the posterior Wnt-Cdx module. Despite the absence of RA signaling, Wnt signaling was elevated in both *Eomes* and *Tbxt* null mutants, correlating with *Wnt3a* upregulation in *Eomes^-/-^* and sustained *Wnt3* expression in *Tbxt^-/-^*. In both the cases, a Cdx2+ posterior growth zone was formed. We propose that deregulation of Wnt signaling in the mutants causes stabilization of the Wnt-Cdx module resulting in posterior fate specification. These findings underscore the central role of Wnt signaling, and suggest that RA functions by gating the Wnt signaling in A-P developmental progression.

RA signaling has a conserved role in regulating the collinear expression of *Hox* genes across Chordata (Schubert et al., 2006). RA signaling activates Cdx factors (Amblard et al., 2025; Houle et al., 2000), implicating RA in the activation of mid-cluster Hox genes via Cdx factors (Deschamps & van Nes, 2005). Moreover, several studies have demonstrated that RA signalling regulates patterning of the posterior body (Abu-Abed et al., 2001; Hennessy et al., 2023; Iulianella et al., 1999; Kessel & Gruss, 1991). Thus, the role of RA in patterning the posterior body following the transition to posterior program is well established. In this study, we have demonstrated that the preclusion of RA signaling is essential for anterior identity specification. In this context, the importance of RA signaling is underscored by the striking phenotype observed in *Cyp26a1/2/3^-/-^* triple mutant embryos. Failure to restrict the influence of maternal RA in early streak mouse embryos leads to ectopic *Tbxt* expression in the anterior region. This has been interpreted as axis duplication (Uehara et al., 2009). Given our findings, we reinterpret the *Cyp26a^-/-^*phenotype as also reflecting the ‘posteriorization’ of anterior mesoderm. Taking this together with our findings, we conclude that the inhibition of RA signaling is essential for safeguarding anterior development from inappropriate posterior differentiation. We propose that the regulation of RA signaling, from its suppression to activation during the early to late phases of gastrulation, is key to regulating the orderly A-P developmental transition. Addressing how RA signalling itself is regulated, in the switch from an RA-precluded to an RA-active state, will further elucidate the anterior-to-posterior developmental transition.

The Wnt and Nodal pathways play a fundamentally conserved role in A-P axis patterning (Arnold & Robertson, 2009; Hikasa & Sokol, 2013; Robertson, 2014; Tian & Meng, 2006). The RA signaling pathway is similarly conserved across eumetazoans (Rhinn & Dollé, 2012). However, RA signaling appears to have been co-opted for patterning the overall A-P body axis later in evolutionary history, suggesting that it is a chordate innovation (Cañestro et al., 2006; Marlétaz et al., 2006). We have demonstrated that Wnt signaling is central and that RA signaling gates Wnt signaling to regulate the axial mesoderm patterning. Taking this together with the later recruitment of RA signaling in phylogeny, we propose that RA signaling functions to reinforce the core Nodal/Wnt mechanism regulating the orderly developmental transition. With the complexity of the animal body increasing through evolution, the co-option of RA signaling could have reinforced the A-P patterning mechanisms by making it robust and resilient to noise.

## Material and methods

### Lead contact

Further information and requests for resources and reagents should be directed to and will be fulfilled by the Lead Contact, [Ramkumar Sambasivan] ().

## Materials availability

Cell lines generated in this study are available upon request.

## Data and code availability

Code for the single cell analysis can be found in (https://github.com/gatocor/project-vitamin-A)

## Experimental model and subject details Mouse embryonic stem cells

All cell lines used are listed in methods table 1 below. Gastruloids were generated from E14Tg2a and E14Tg2a-GFP mouse embryonic stem cells (mESCs). We have also used signaling pathway reporter lines such as Tcf/Lef-mCherry and AR8-mCherry, and RARE-d2-eGFP. *Eomes^-/-^* and *Tbxt^-/-^* mESCs were generated in-house using the E14Tg2a line. *Eomes^-/-^-GFP* and *Tbxt^-/-^-tdTomato* were used for loss of function studies.

**Methods table 1.**
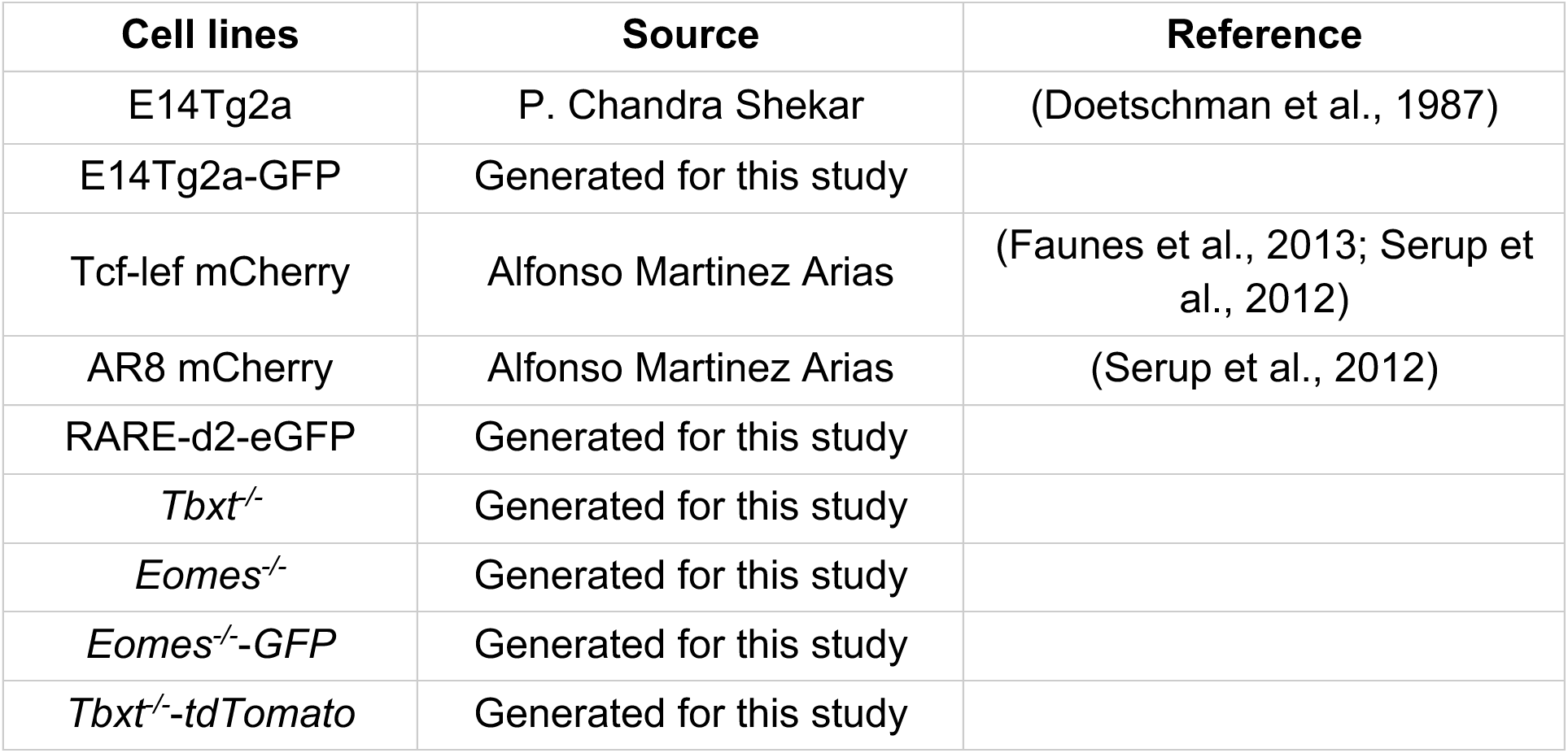

| Cell lines | Source | Reference |
| --- | --- | --- |
| E14Tg2a | P. Chandra Shekar | (Doetschman et al., 1987) |
| E14Tg2a-GFP | Generated for this study |  |
| Tcf-lef mCherry | Alfonso Martinez Arias | (Faunes et al., 2013; Serup et al., 2012) |
| AR8 mCherry | Alfonso Martinez Arias | (Serup et al., 2012) |
| RARE-d2-eGFP | Generated for this study |  |
| *Tbxt*<sup>-/-</sup> | Generated for this study |  |
| *Eomes*<sup>-/-</sup> | Generated for this study |  |
| *Eomes*<sup>-/-</sup>-GFP | Generated for this study |  |
| *Tbxt*<sup>-/-</sup>-tdTomato | Generated for this study |  |

## METHOD DETAILS

### Cell culture

Mouse embryonic stem cells (mESCs) were maintained at 37°C and 5% CO2 on 0.1% gelatin-coated 35-mm tissue culture dishes. Glasgow Minimum Essential Medium (GMEM) supplemented with 10% fetal bovine serum (FBS), 1× non-essential amino acids, 2mM GlutaMAX, 1× penicillin–streptomycin, 0.1mM β-mercaptoethanol, and 1000U/mL leukemia inhibitory factor (LIF; ESGRO) was used as the growth medium. Cells were passaged every other day at a confluency of ≤60%. For passaging, the medium was removed, the cells were washed with 1mL of PBS, and then dissociated with 400µL of TrypLE for 3.5 min at 37 °C. The reaction was quenched with 1.6mL of growth medium, and the cells were triturated 15–20 times to obtain a single-cell suspension. One-tenth of the suspension was transferred to a fresh gelatin-coated dish containing 1.8mL of growth medium. The references for media and culture reagents and plasticware are listed in methods tables 2 and 3.

**Methods table 2.**

| mES growth media | Concentration | Catalogue number | Source |
| --- | --- | --- | --- |
| GMEM | Make up to the required volume | 11710035 | Gibco |
| Fetal bovine serum | 10% | RM9952 OR RM10686 | Gibco |
| MEM non-essential amino acid | 1X | 11140-050 | Gibco |
| Glutamax (100X) | 1X | 35050-061 | Gibco |
| Penicillin/ Streptomycin (100X) | 1X | 15070063 | Gibco |
| β-mercaptoethanol | 100 μM | 21985023 | Gibco |
| Leukemia inhibitory factor (LIF) | 10<sup>3</sup> units (1:10000 dilution) | ESG1107 | Merck |

**Methods table 3.**

| Cell culture | Catalogue number | Source |
| --- | --- | --- |
| PBS, pH 7.4 | 10010023 | Gibco |
| TrypLE™ Express Enzyme (1X), Phenol Red | 12605010 | Gibco |
| Gelatin | TC041 - 100G | Himedia |
| 35-mm tissue culture dish | 353001 | Corning |
| 60mm tissue culture dish | 353002 | Corning |
| 96-well U bottom dish,<br>non-treated | 34296 | Genetix |
| 6-well low attachment<br>dish | 3471 | Corning |

### Gastruloid culture

The mESCs were preconditioned by subculturing in 35-mm dishes on growth medium supplemented with 1μM PD0325901 (MEK inhibitor) and 3μM CHIR99021 (GSK3 inhibitor) for 24h prior to gastruloid formation. Following the 2i pre-treatment, mESCs were washed twice with 1mL PBS and dissociated with TrypLE (similar to the cell passage protocol). The single-cell suspension was transferred to a 15mL tube and centrifuged at 1000 rpm for 3 min in a swing-bucket centrifuge. The supernatant was removed, and the cells were washed twice with 2mL PBS. Pelleted cells were resuspended in N2B27 medium, and viable cell numbers were determined using a hemocytometer. N2B27 medium was prepared by combining 24.1mL DMEM/F12, 24.1mL Neurobasal medium, 250μL N2 supplement, 500μL B27 supplement, 500μL L-glutamine, 500μL penicillin–streptomycin, and 50μL β-mercaptoethanol to a final volume of 50mL. For posterior mesoderm (PM) gastruloids, B27 supplement with vitamin A was used, whereas for anterior mesoderm (AM) gastruloids, B27 without vitamin A was used.

Aggregates were generated by seeding 300 cells per well in 40μL N2B27 medium in non-tissue-culture-treated U-bottom 96-well plates. After 48h, gastruloids were induced with 3μM CHIR99021 in 150μL N2B27. At 72h and 96h, the spent medium was replaced with 150μL of fresh N2B27. For endpoint analysis, gastruloids were collected at 120h. The references for media and other reagents are listed in methods tables 4 and 5.

**Methods table 4.**

| N2B27 media | Working Concentration | Catalogue number | Source |
| --- | --- | --- | --- |
| Neurobasal medium | 48.20% | 21103049 | Gibco |
| DMEM-F12 (HEPES) | 48.20% | 11330032 | Gibco |
| N2 Supplement (100X) | 1X | 17502048 | Gibco |
| B27 with vitamin A (50X) | 1X | 17504044 | Gibco |
| B27 without vitamin A (50X) | 1X | 12587010 | Gibco |
| L-glutamine (100X) | 1X | 25030081 | Gibco |
| Penicillin/ Streptomycin (100X) | 1X | 15070063 | Gibco |
| β-mercaptoethanol | 100μM | 21985023 | Gibco |

**Methods table 5.**

| **Growth factors and inhibitors** | **Concentration** | **Catalogue number** | **Source** |
| --- | --- | --- | --- |
| CHIR99021 (GSK3 inhibitor) | 3μM | 4423/10 | Tocris |
| CHIR99021 (GSK3 inhibitor) | 3μM | SML1046-25MG | Merck |
| PD0325901 (MEK inhibitor) | 1μM | 100-0248 | Stemcell Technologies |
| Activin A | 100ng/ml | 338-AC-050 | R&D Systems |
| Xav939 | 5μM | X3004-5MG | Sigma |
| Fgf10 | 50ng/ml | 345-FG-025 | R&D Systems |
| IGF1 | 2ng/ml | 250-19-10 | Peprtech |

### Cardiac organoid culture

After 120h, the AM gastruloids were transferred to a 6-well low attachment plate and kept on a shaker at 40-60rpm and continued in the gastruloid culture medium supplemented with Fgf10 (50ng/ml) and IGF-1 (2ng/ml). For endpoint analysis, gastruloids were collected at 192h.

### Immunostaining and imaging

Gastruloids were collected into individual wells of a 48-well plate or 24-well plate, rinsed twice with PBS to remove residual medium, and fixed in 4% paraformaldehyde (PFA) for 2h at 4 °C. Following fixation, gastruloids were washed three times with PBS (20 min each at room temperature) and stored at 4 °C until further processing. Gastruloids were permeabilized and blocked in PBS-FT (PBS containing 10% FBS and 0.25% Triton X-100) for 1h at room temperature. All staining steps were performed on an orbital shaker. Primary antibodies were diluted in PBS-FT at the indicated concentrations, and gastruloids were incubated for 24h at 4 °C. Samples were then washed three times with PBS-FT (20 min each) and incubated with secondary antibodies, also diluted in PBS-FT, for 24h at 4 °C. After staining, gastruloids were washed three additional times with PBS-FT (20 min each), rinsed with PBS, and stored at 4 °C until imaging.

Gastruloids were mounted in 100% glycerol on coverslip-bottom 35-mm dishes. Confocal images were acquired using an Olympus FV3000 microscope with 10× or 20× objectives. Z-stacks were captured using system-optimized step sizes.

Post-acquisition image processing was performed in Fiji (ImageJ). Fluorescence channels were split, and maximum intensity projections were generated from the acquired Z-stacks for preparing the images for the figures. For quantification of fluorescence intensity along the anterior-posterior (A-P) axis, a midline was drawn along the length of each gastruloid using the line tool and fitted to a spline. Intensity values along the midline were extracted using the “Plot Profile” function (Cmd+K), exported to Excel, and plotted in R. For elongation index quantification, the major axis was defined as the length of the midline, and the minor axis was measured as the diameter of the largest inscribed circle within the gastruloid.

The primary antibodies used are listed in table 6.

**Methods table 6.**
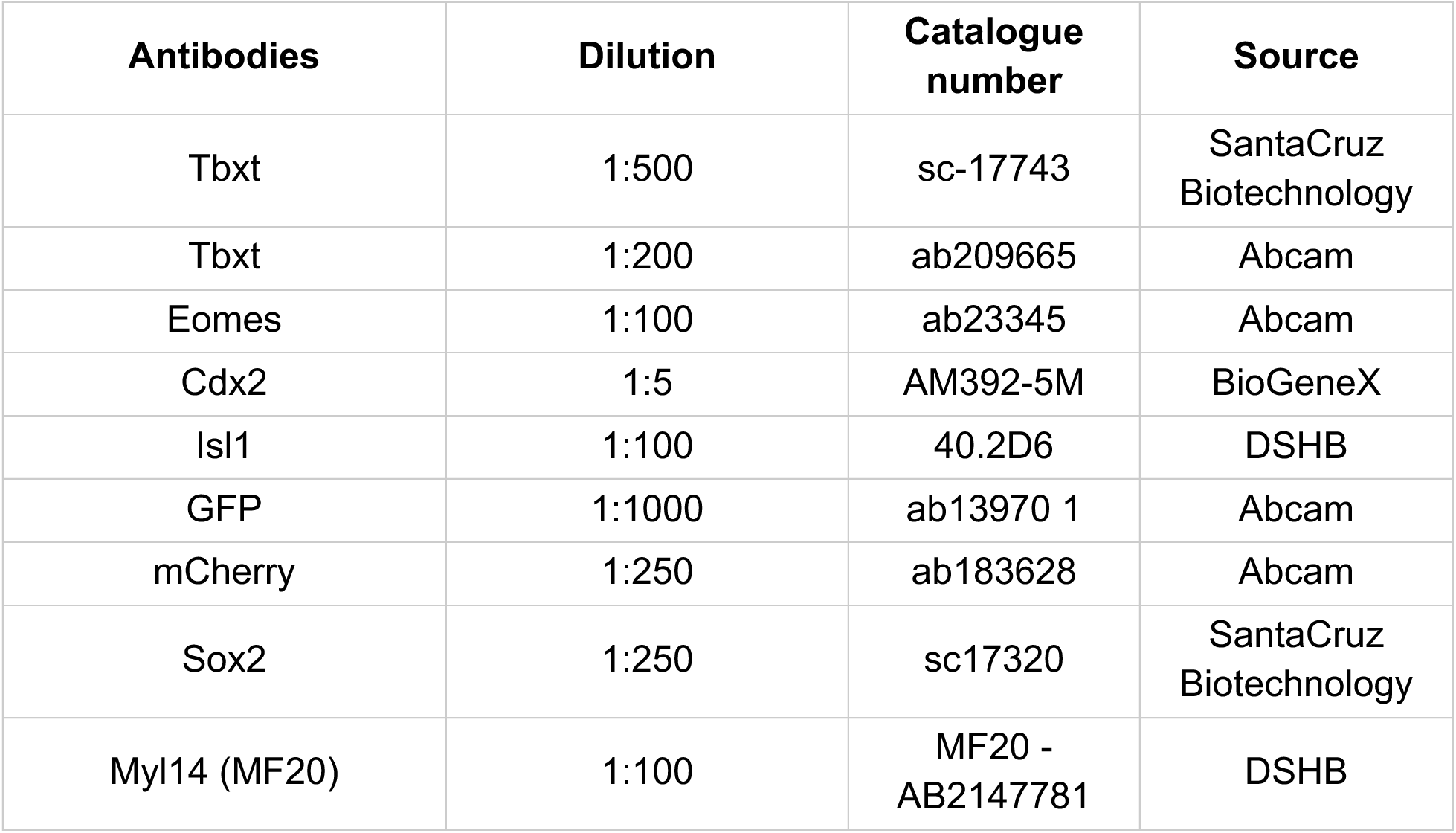

### *in situ* hybridization (ISH) Probe synthesis

Antisense RNA probes were generated either from linearized plasmid templates or from PCR-amplified fragments. For plasmid-based templates, vectors containing the desired insert were linearized, and *in vitro* transcription was performed using 1µg of linearized DNA with the DIG RNA Labeling Kit (Roche, Cat. No. 11175025910), according to the manufacturer’s instructions. The reaction mixture contained 1mM each of ATP, CTP, and GTP, 0.65mM UTP, and 0.35mM DIG-11-UTP and was incubated for 2h at 37°C. Template DNA was then digested with DNase I, and the reaction products were purified using Bio-Rad Micro Bio-Spin 30 chromatography columns. Probe integrity was verified by agarose gel electrophoresis, and concentrations were determined by a NanoDrop spectrophotometer.

For probes generated from PCR-amplified DNA templates, cDNA was synthesized from mouse embryo RNA corresponding to a developmental stage known to express the gene of interest, and cDNA was synthesized. Gene-specific primers were designed to amplify the desired fragment, with a T7 or T3 RNA polymerase binding site appended to the 5′ end of the forward primer. The purified PCR amplicon was then used directly as a template for *in vitro* transcription, as described above. Aliquots of the purified probes (typically ∼200ng per reaction were stored at-20°C until use.

### Whole-mount ISH on gastruloids

Gastruloids were collected in 4-well dishes. Media was removed, and gastruloids were washed with PBS before fixing in 4% paraformaldehyde (PFA) at 4°C overnight. Three washes in PTW (PBS + 0.1% Tween-20) were given to remove PFA. Samples were dehydrated through a methanol/PTW series (25%, 50%, 75%, 100%) and stored in 100% methanol at –20°C. On the day of performing in-situ hybridization, samples were rehydrated in methanol/PTW series (75%, 50% 20%, 0%). No Proteinase K treatment was done prior to hybridization. Prehybridization was performed for 1h at 70°C in hybridization mix (50% formamide, 1.3× SSC (pH 5), 5mM EDTA, 0.5% CHAPS, 0.2% Tween-20, 100µg/mL heparin, and 50µg/mL yeast RNA). Hybridization was carried out overnight at 70°C with ∼100ng of DIG-labeled RNA probe in a pre-warmed hybridization mix. Post-hybridization washes were performed at 70°C (2 × 30 min in hybridization mix, followed by 1 × 20 min in a 1:1 hybridization mix/MABT solution). Gastruloids were then washed three times in MABT (Maleic acid buffer + 0.1% Tween-20) and blocked for 1h in MABT containing 2% Roche blocking reagent and 20% heat-inactivated goat serum. Anti-DIG-AP antibody (Roche) was added at a 1:2000 dilution in the blocking buffer and incubated overnight at 4°C. Following three washes in MABT and a 24h room-temperature wash, gastruloids were rinsed twice in NTMT buffer and stained in BM Purple (Roche) at room temperature until color development was complete. Staining reactions were stopped by washing three times in PTW. Gastruloids were post-fixed overnight in 4% PFA at 4°C and stored long-term in 0.5% PFA at 4°C. Stained gastruloids were imaged on a 35-mm dish in PBS using a stereo zoom microscope. The reagent list is provided in table 7.

**Methods table 7.**

| Components of ISH Hybridization Buffer | Concentration | Stock Concentration | Catalogue number | Source |
| --- | --- | --- | --- | --- |
| Formamide | 50% | - | 47671-1L-F | Sigma |
| SSC (pH 5) | 1.3x | 20x | S6639-1L | Sigma |
| EDTA | 5mM | 0.5M, pH 8.0 | 15575020 | Gibco |
| Yeast RNA | 50µg/ mL | 10mg /mL | 1010922300<br>1 | Roche |
| CHAPS | 0.5% | 10% | C3023-5G | Sigma |
| Tween-20 | 0.2% | 10% | P1379 | Sigma |
| Heparin | 100µg/ mL | 50mg/ mL | H3149 | Sigma |

### Hybridization Chain Reaction

(HCR; RNA Fluorescence *in situ* Hybridization):

HCR RNA-FISH was performed using the Molecular Instruments HCR v3.0 kit (Molecular Instruments, Los Angeles, CA) according to the manufacturer’s instructions, with minor modifications optimized for gastruloids and mouse embryos. Briefly, samples were fixed, dehydrated, and rehydrated similar to ISH as mentioned above. Prior to hybridization, samples were equilibrated in HCR hybridization buffer (Molecular Instruments). The pre-hybridization solution was then removed and replaced with fresh hybridization buffer containing probe sets targeting the genes of interest (typically at a final probe concentration of 4nM) and incubated overnight at 37°C. Following hybridization, excess probes were removed by a series of washes in pre-warmed HCR wash buffer and 5× SSCT (saline-sodium citrate with 0.1% Tween-20), as per the Molecular Instruments protocol.

For signal amplification, samples were equilibrated in amplification buffer and then incubated overnight at room temperature in amplification buffer containing pre-annealed fluorescently labeled HCR hairpins (typically 60nM each). Prior to use, hairpins were snap-cooled by heating to 95°C for 90s followed by slow cooling to room temperature for 30 min in the dark.

After amplification, samples were washed extensively in 5× SSCT to remove unbound hairpins and then counterstained with Hoechst (1µg/ml) for 48h at 4°C. Imaging was performed using a confocal laser scanning microscope (Olympus FV3000) with identical acquisition settings across conditions.

### Single-cell RNA sequencing

Single-cell suspensions were prepared from gastruloids derived from wild type (n = 3 biological replicates), *Eomes* mutant (n = 2), and *Tbxt* mutant (n = 2) lines. For each experiment, ∼96 gastruloids were collected in a 35-mm dish and washed once with PBS. Gastruloids were dissociated with 700μL TrypLE Express for 5 min at 37°C and quenched with 3.2mL FBS-containing growth medium. The suspension was gently triturated to obtain a single-cell suspension and transferred to a 15mL tube. The suspension was diluted with 10mL PBS, and cells were pelleted by centrifugation (1000 rpm, 3 min). The pellet was resuspended in PBS supplemented with 1% BSA, and cell concentration and viability were determined using a hemocytometer. The viability was ∼96% in the cell suspension.

Single-cell RNA sequencing libraries were prepared using the Chromium Next GEM Single Cell 3′ Library & Gel Bead Kit v3.1 (10x Genomics) according to the manufacturer’s instructions. For each condition, approximately 8,000 dissociated cells were loaded per channel. Libraries were sequenced on an Illumina NovaSeq 6000 (paired-end, 28 × 91 bp), targeting ∼50,000 reads per cell. The mean number of reads per cell ranged from 20,000 to 50,000 across samples. Raw data were processed using Cell Ranger v9.0.0 (10x Genomics) with the mm10 reference genome (refdata-gex-mm10-2020-A).

All analyses were performed using Scanpy (Wolf et al., 2018) unless otherwise stated.

Quality control was performed independently for each condition and replicate. Standard QC metrics were computed, including the total number of transcripts per cell, the number of detected genes, and the fraction of mitochondrial reads. Thresholds for filtering low-quality cells were determined heuristically from the distributions of these metrics to remove cells with low RNA content, stressed or dying cells, empty droplets, and potential doublets. Doublet detection was refined using Scrublet (Wolock et al., 2019). The doublet score threshold was chosen heuristically based on UMAP representations to exclude regions enriched for high doublet scores while maintaining biological integrity across batches. Outlier cells with abnormally high mean distances from their k-nearest neighbors were also excluded. Inclusion or exclusion of these cells did not alter the number or identity of clusters, confirming that no biologically relevant populations were removed.

Count matrices were normalized such that all cells had the same total number of counts, corresponding to the mean library size across the dataset. Normalized counts were log-transformed using the log1p function. The resulting log-normalized matrices were used for all downstream analyses, including dimensionality reduction and clustering.

Highly variable genes (HVGs) were identified using Scanpy’s default parameters, retaining the top 2,000 genes. Principal component analysis (PCA) was performed on the HVG expression matrix, and the number of principal components retained was determined using the elbow heuristic based on the explained variance ratio.

Analyses involving multiple batches or experimental conditions were batch-corrected using Harmony (Korsunsky et al., 2019) applied to the PCA representation. Cell cycle phase was annotated using curated gene lists (Tirosh et al., 2016), and, where necessary, a second Harmony integration was performed to mitigate residual cell cycle–related effects.

A k-nearest neighbor (KNN) graph (k = 15) was constructed from the batch-and cell-cycle–corrected PCA space. Two-dimensional representations were generated using UMAP for the visualization of cell population structure.

Clusters were identified using the Leiden algorithm (Traag et al., 2019) on the KNN graph. Differential expression analysis was performed using the Wilcoxon rank-sum test with Hochberg correction for multiple testing. Cluster annotation was guided by differentially expressed genes and curated marker gene lists, visualized on UMAPs and summarized in dot plots. P-values from differential expression analysis were interpreted as indicative rather than definitive due to the inherent dependence between clustering and differential expression testing.

Datasets were projected onto reference mouse embryo and gastruloid single-cell atlases using a custom implementation of scmap (Kiselev et al., 2018). Gene identifier discrepancies (e.g., Ensembl IDs vs. common gene symbols) were resolved using mygene.info (Wu et al., 2013), and duplicated mappings were removed to ensure consistent one-to-one correspondence prior to projection.

## Real-time quantitative PCR

### Sample Collection and RNA Extraction

The number of gastruloids harvested per time point was as follows: 48h, 96 gastruloids; 60h, 96 gastruloids; 72h, 72 gastruloids; 96h, 72 gastruloids; and 120h, 48 gastruloids. Gastruloids were rinsed once with PBS and lysed in 350 µL of lysis buffer from the HiGenoMB HiPurA Total RNA Miniprep Purification Kit (HiMedia) supplemented with 3.5µL of 13.4M β-mercaptoethanol. Total RNA was extracted following the manufacturer’s instructions. RNA yield ranged from 80 to 350ng/µL. For each sample, 1µg of total RNA was reverse transcribed using the PrimeScript First Strand cDNA Synthesis Kit (Takara, Cat. No. 6110A) according to the manufacturer’s protocol. qPCR reactions were set up in MicroAmp Optical 384-well reaction plates (Applied Biosystems, Cat. No. 4309849). Reactions were prepared using Maxima SYBR Green/ROX qPCR Master Mix (Thermo Fisher Scientific) and gene-specific primers. Ct values were obtained for each gene and normalized to Actin (ΔCt). For temporal comparisons, relative expression for each gene was normalized to the corresponding 48-h time point within each condition (ΔΔCt).

Statistical analyses were performed in GraphPad Prism. A two-way ordinary ANOVA was performed with time as the row factor (changes in gene expression across time), condition as the column factor (comparing global between PM and AM), and their interaction (to determine whether temporal expression patterns differ between conditions). It was followed by Sidak’s post-hoc test to identify significantly different time points between conditions. To evaluate changes within a condition relative to 48h, ΔCt values were compared using a two-tailed paired t-test. Line graphs represent mean ± SD with data points connected to illustrate temporal trends.

The list of primers used for RT-qPCR are provided in table 8.

**Methods table 8.**
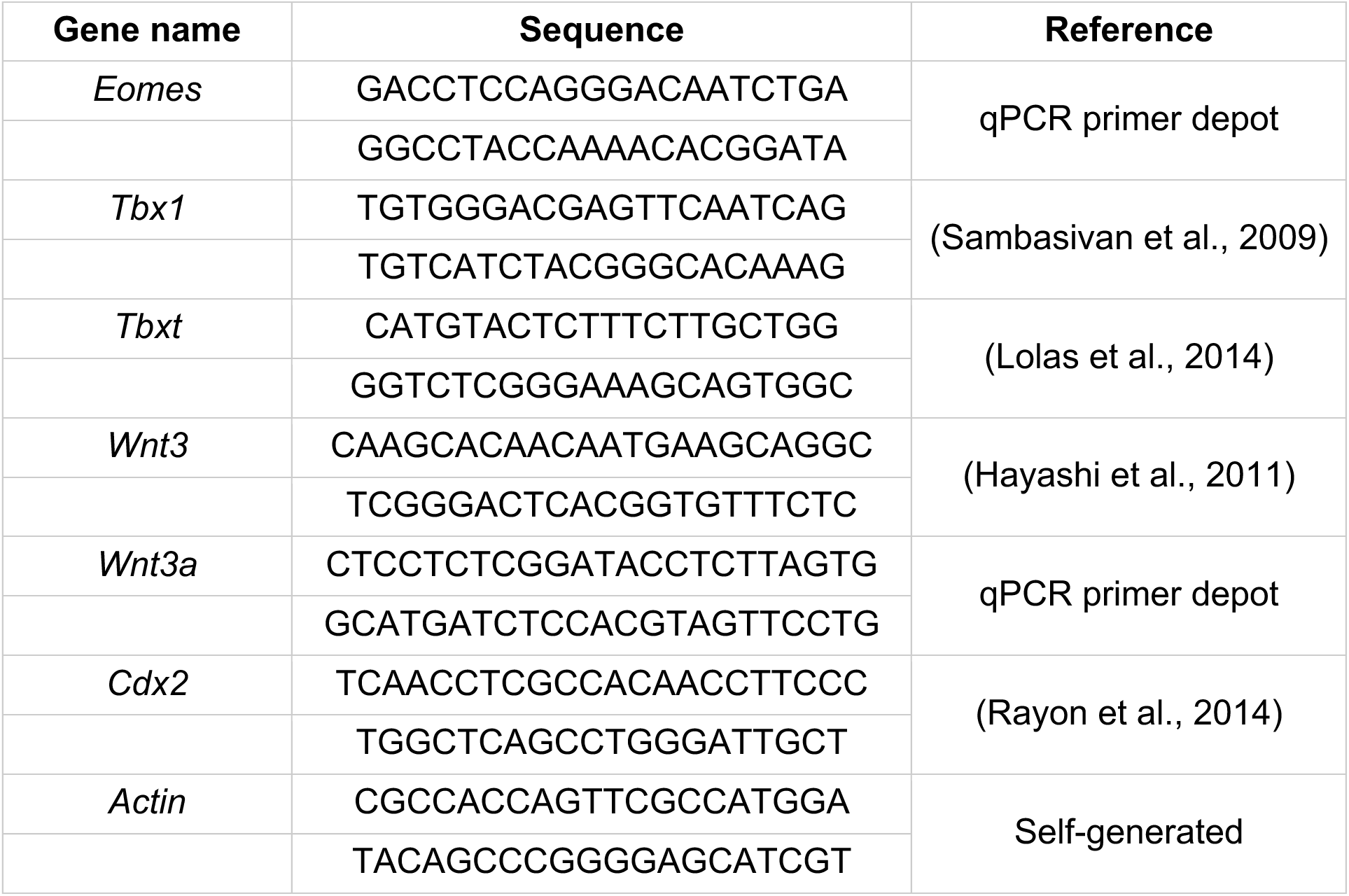

### Flow cytometry analysis

Single-cell suspensions were generated from 24 120h gastruloids per condition, following the dissociation protocol described in the *single-cell RNA sequencing* section. After centrifugation, cell pellets were resuspended in PBS and analyzed on a Beckman Coulter CytoFLEX-S flow cytometer.

During post-acquisition analysis, debris was excluded by gating on FSC-A versus SSC-A. To define the negative population, the unlabelled E14Tg2a cell line was used as a control. For the *Tcf/Lef:mCherry* and *AR8:mCherry* reporter experiments comparing PM and AM gastruloids, a fluorescence threshold was set such that 95% of the negative-control events fell within the negative gate. Median fluorescence intensity was then calculated using only the events above this threshold.

For the chimeric gastruloid experiments, a single universal threshold could not be applied because the samples contained mixtures of multiple cell lines, each exhibiting different background fluorescence levels. As expected for chimeric samples, all conditions contained both reporter-negative and reporter-positive cells, resulting in a bimodal distribution of fluorescence intensities. Therefore, for each condition, the local minimum between the two peaks was used as the cutoff to separate the negative and positive populations. Median fluorescence intensity was subsequently calculated using only the reporter-positive cells.

## Acknowledgements

We thank Prof. Alfonso Martinez Arias and Dr. Andre Dias for their insightful discussions and valuable feedback throughout this work. We are grateful to Mr. Prahlad Kashyap for assistance with line-scan quantitations. We acknowledge Dr. Awadhesh Pandit, Mr. Santosh Srirangam, Mr. Lakshminarayanan C.P. and Dr. H. Krishnamurthy for their support with single-cell sequencing experiments. We also thank Dr. Rajesh Ladher, Dr. Praveen Vemula, Dr. Dasaradhi Palakodeti and Dr. Dhandapani Perundurai for hosting S.K.Y. and A.B. and facilitating the single-cell sequencing experiments in Bangalore.

This research was supported by the Department of Biotechnology, Government of India (grants BT/PR39986/MED/97/479/2020 and BT/PR45132/MED/97/621/2022) and by IISER Tirupati core funds to R.S. A.B. is supported by a UGC-JRF/SRF fellowship, S.K.Y. by an IISER Tirupati Integrated PhD fellowship, and B.V. was supported by DBT-JRF/SRF fellowships and an InStem fellowship.

## Author Contributions

Conceptualization, R.S., A.B., and S.K.Y.; Methodology, A.B., S.K.Y., B.V., and R.S.; Investigation, A.B., S.K.Y., and B.V.; Formal Analysis, R.S., A.B., S.K.Y., and G.T.C.; Visualization, G.T.C.; Resources, B.V., A.B, S.K.Y., and R.S.; Supervision, R.S.;

Writing – Original Draft, R.S., A.B., and S.K.Y.; Writing – Review & Editing, R.S., A.B., and S.K.Y.

## Notes

### Competing Interest Statement

The authors have declared no competing interest.

### Summary of Updates

The introduction and discussion sections have been revised to be more focused and easier to read. New results have been included in some of the figures.

## References

Abu-Abed, S., Dollé, P., Metzger, D., Beckett, B., Chambon, P., & Petkovich, M. (2001). The retinoic acid-metabolizing enzyme, CYP26A1, is essential for normal hindbrain patterning, vertebral identity, and development of posterior structures. Genes & Development, 15(2), 226. 10.1101/GAD.855001

Amblard, I., Baranasic, D., Xie, S. Q., Moyon, B., Percharde, M., Lenhard, B., & Metzis, V. (2025). A dual enhancer-attenuator element ensures transient Cdx2 expression during mouse posterior body formation. Developmental Cell, 60(18). 10.1016/j.devcel.2025.06.006

Amin, S., Neijts, R., Simmini, S., Van Rooijen, C., Tan, S. C., Kester, L., Van Oudenaarden, A., Creyghton, M. P., & Deschamps, J. (2016). Cdx and T Brachyury Co-activate Growth Signaling in the Embryonic Axial Progenitor Niche. Cell Reports, 17(12), 3165–3177. 10.1016/j.celrep.2016.11.069

Ang, H. L., Deltour, L., Hayamizu, T. F., Žgombić-Knight, M., & Duester, G. (1996). Retinoic acid synthesis in mouse embryos during gastrulation and craniofacial development linked to class IV alcohol dehydrogenase gene expression. Journal of Biological Chemistry, 271(16), 9526–9534. 10.1074/JBC.271.16.9526

Arnold, S. J., Hofmann, U. K., Bikoff, E. K., & Robertson, E. J. (2008). Pivotal roles for eomesodermin during axis formation,epithelium-to-mesenchyme transition and endoderm specification in the mouse. Development, 135(3), 501–511. 10.1242/dev.014357

Arnold, S. J., & Robertson, E. J. (2009). Making a commitment: cell lineage allocation and axis patterning in the early mouse embryo. Nature Reviews Molecular Cell Biology 2009 10:2, 10(2), 91–103. 10.1038/nrm2618

Beccari, L., Moris, N., Girgin, M., Turner, D. A., Baillie-Johnson, P., Cossy, A. C., Lutolf, M. P., Duboule, D., & Arias, A. M. (2018). Multi-axial self-organization properties of mouse embryonic stem cells into gastruloids. Nature 2018 562:7726, 562(7726), 272–276. 10.1038/s41586-018-0578-0

Cañestro, C., Postlethwait, J. H., Gonzàlez-Duarte, R., & Albalat, R. (2006). Is retinoic acid genetic machinery a chordate innovation? Evolution and Development, 8(5), 394–406. 10.1111/J.1525-142X.2006.00113.X

Chapman, D. L., & Papaioannou, V. E. (1998). Three neural tubes in mouse embryos with mutations in the T-box gene Tbx6. Nature 1998 391:6668, 391(6668), 695–697. 10.1038/35624

Chawengsaksophak, K., De Graaff, W., Rossant, J., Deschamps, J., & Beck, F. (2004). Cdx2 is essential for axial elongation in mouse development. Proceedings of the National Academy of Sciences of the United States of America, 101(20), 7641–7645. 10.1073/PNAS.0401654101;PAGEGROUP:STRING:PUBLICATION

Ciruna, B., & Rossant, J. (2001). FGF Signaling Regulates Mesoderm Cell Fate Specification and Morphogenetic Movement at the Primitive Streak. Developmental Cell, 1(1), 37–49. 10.1016/S1534-5807(01)00017-X

Costello, I., Pimeisl, I. M., Dräger, S., Bikoff, E. K., Robertson, E. J., & Arnold, S. J. (2011). The T-box transcription factor Eomesodermin acts upstream of Mesp1 to specify cardiac mesoderm during mouse gastrulation. Nature Cell Biology 2011 13:9, 13(9), 1084–1091. 10.1038/ncb2304

Deschamps, J., & van Nes, J. (2005). Developmental regulation of the Hox genes during axial morphogenesis in the mouse. Development, 132(13), 2931–2942. 10.1242/DEV.01897

Dias, A., Pascual-Mas, P., Robertson, G., Torregrosa-Cortés, G., Stelloo, S., Casaní-Galdón, P., Babin, S., Romaniuk, Y., Mayran, A., Wehmeyer, A. E., Garcia-Ojalvo, J., McNamara, H. M., Vermeulen, M., Arnold, S. J., & Arias, A. M. (2025). Opposing Nodal and Wnt signalling activities govern the emergence of the mammalian body plan. BioRxiv, 2025.01.11.632562. 10.1101/2025.01.11.632562

Doetschman, T., Gregg, R. G., Maeda, N., Hooper, M. L., Melton, D. W., Thompson, S., & Smithies, O. (1987). Targetted correction of a mutant HPRT gene in mouse embryonic stem cells. Nature, 330(6148), 576–578. 10.1038/330576A0;KWRD

Durston, A. J., Timmermans, J. P. M., Hage, W. J., Hendriks, H. F. J., De Vries, N. J., Heideveld, M., & Nieuwkoop, P. D. (1989). Retinoic acid causes an anteroposterior transformation in the developing central nervous system. Nature 1989 340:6229, 340(6229), 140–144. 10.1038/340140a0

Ee, L. sha, Medina-Cano, D., Goetzler, E., Uyehara, C., Schwarz, C., Salataj, E., Madhuranath, S., Evans, T., Hadjantonakis, A. K., Apostolou, E., Polyzos, A., Vierbuchen, T., & Stadtfeld, M. (2025). Enhancer remodeling by OTX2 directs specification and patterning of mammalian definitive endoderm. Developmental Cell, 0(0). 10.1016/j.devcel.2025.07.020

Evans, A. L., Faial, T., Gilchrist, M. J., Down, T., Vallier, L., Pedersen, R. A., Wardle, F. C., & Smith, J. C. (2012). Genomic targets of Brachyury (T) in differentiating mouse embryonic stem cells. PloS One, 7(3). 10.1371/JOURNAL.PONE.0033346

Faunes, F., Hayward, P., Descalzo, S. M., Chatterjee, S. S., Balayo, T., Trott, J., Christoforou, A., Ferrer-Vaquer, A., Hadjantonakis, A. K., Dasgupta, R., & Arias, A. M. (2013). A membrane-associated β-catenin/Oct4 complex correlates with ground-state pluripotency in mouse embryonic stem cells. Development (Cambridge), 140(6), 1171–1183. 10.1242/DEV.085654/VIDEO-3

Ferrer-Vaquer, A., Piliszek, A., Tian, G., Aho, R. J., Dufort, D., & Hadjantonakis, A. K. (2010). A sensitive and bright single-cell resolution live imaging reporter of Wnt/ß-catenin signaling in the mouse. BMC Developmental Biology 2010 10:1, 10(1), 121-. 10.1186/1471-213X-10-121

Gouti, M., Tsakiridis, A., Wymeersch, F. J., Huang, Y., Kleinjung, J., Wilson, V., & Briscoe, J. (2014). In Vitro Generation of Neuromesodermal Progenitors Reveals Distinct Roles for Wnt Signalling in the Specification of Spinal Cord and Paraxial Mesoderm Identity. PLOS Biology, 12(8), e1001937. 10.1371/JOURNAL.PBIO.1001937

Hayashi, K., Ohta, H., Kurimoto, K., Aramaki, S., & Saitou, M. (2011). Reconstitution of the mouse germ cell specification pathway in culture by pluripotent stem cells. Cell, 146(4), 519–532. 10.1016/j.cell.2011.06.052

Hennessy, M. J., Fulton, T., Turner, D. A., & Steventon, B. (2023). Negative feedback on Retinoic Acid by Brachyury guides gastruloid symmetry-breaking. BioRxiv, 2023.06.02.543388. 10.1101/2023.06.02.543388

Herrmann, B. G. (1991). Expression pattern of the Brachyury gene in whole-mount TWIS/TWIS mutant embryos. Development, 113(3), 913–917. 10.1242/DEV.113.3.913

Hikasa, H., & Sokol, S. Y. (2013). Wnt Signaling in Vertebrate Axis Specification. Cold Spring Harbor Perspectives in Biology, 5(1), a007955. 10.1101/CSHPERSPECT.A007955

Ikeya, M., & Takada, S. (2001). Wnt-3a is required for somite specification along the anteroposterior axis of the mouse embryo and for regulation of cdx-1 expression. Mechanisms of Development, 103(1–2), 27–33. 10.1016/s0925-4773(01)00338-0

Iulianella, A., Beckett, B., Petkovich, M., & Lohnes, D. (1999). A Molecular Basis for Retinoic Acid-Induced Axial Truncation. Developmental Biology, 205(1), 33–48. 10.1006/DBIO.1998.9110

Javali, A., Misra, A., Leonavicius, K., Acharyya, D., Vyas, B., & Sambasivan, R. (2017). Co-expression of Tbx6 and Sox2 identifies a novel transient neuromesoderm progenitor cell state. Development (Cambridge), 144(24), 4522–4529. 10.1242/DEV.153262/264518/AM/CO-EXPRESSION-OF-TBX6-AND-SOX2-IDENTIFIES-A-NOVEL

Jho, E., Zhang, T., Domon, C., Joo, C.-K., Freund, J.-N., & Costantini, F. (2002). Wnt/β-Catenin/Tcf Signaling Induces the Transcription of Axin2, a Negative Regulator of the Signaling Pathway. Molecular and Cellular Biology, 22(4), 1172–1183. 10.1128/MCB.22.4.1172-1183.2002;PAGE:STRING:ARTICLE/CHAPTER

Kessel, M. (1992). Respecification of vertebral identities by retinoic acid. Development, 115(2), 487–501. 10.1242/DEV.115.2.487

Kessel, M., & Gruss, P. (1991). Homeotic transformations of murine vertebrae and concomitant alteration of Hox codes induced by retinoic acid. Cell, 67(1), 89–104. 10.1016/0092-8674(91)90574-I

Kinder, S. J., Hadjantonakis, A.-K., Quinlan, G. A., Tam, P. P. L., Nagy, A., & Tsang, T. E. (1999). The orderly allocation of mesodermal cells to the extraembryonic structures and the anteroposterior axis during gastrulation of the mouse embryo. Development, 126(21), 4691–4701. 10.1242/dev.126.21.4691

Kiselev, V. Y., Yiu, A., & Hemberg, M. (2018). scmap: projection of single-cell RNA-seq data across data sets. Nature Methods 2018 15:5, 15(5), 359–362. 10.1038/nmeth.4644

Korsunsky, I., Millard, N., Fan, J., Slowikowski, K., Zhang, F., Wei, K., Baglaenko, Y., Brenner, M., Loh, P. ru, & Raychaudhuri, S. (2019). Fast, sensitive and accurate integration of single-cell data with Harmony. Nature Methods 2019 16:12, 16(12), 1289–1296. 10.1038/s41592-019-0619-0

Langston, A. W., & Gudas, L. J. (1994). Retinoic acid and homeobox gene regulation. Current Opinion in Genetics & Development, 4(4), 550–555. 10.1016/0959-437X(94)90071-A

Lawson, K. A., Meneses, J. J., & Pedersen, R. A. (1991). Clonal analysis of epiblast fate during germ layer formation in the mouse embryo. Development, 113(3), 891–911. 10.1242/dev.113.3.891

Lickert, H., Huls, G., Clevers, H., Kemler, R., Domon, C., Duluc, I., Freund, J.-N., Meyer, B. I., & Wehrle, C. (2000). Wnt/(beta)-catenin signaling regulates the expression of the homeobox gene Cdx1 in embryonic intestine. Development, 127(17), 3805–3813. 10.1242/dev.127.17.3805

Lolas, M., Valenzuela, P. D. T., Tjian, R., & Liu, Z. (2014). Charting Brachyury-mediated developmental pathways during early mouse embryogenesis. Proceedings of the National Academy of Sciences of the United States of America, 111(12), 4478–4483. 10.1073/PNAS.1402612111;WEBSITE:WEBSITE:PNAS-SITE;ISSUE:ISSUE:DOI

Marlétaz, F., Holland, L. Z., Laudet, V., & Schubert, M. (2006). Retinoic acid signaling and the evolution of chordates. International Journal of Biological Sciences, 2(2), 38. 10.7150/IJBS.2.38

Martin, B. L., & Kimelman, D. (2008). Regulation of Canonical Wnt Signaling by Brachyury Is Essential for Posterior Mesoderm Formation. Developmental Cell, 15(1), 121–133. 10.1016/j.devcel.2008.04.013

Martin, B. L., & Kimelman, D. (2009). Wnt Signaling and the Evolution of Embryonic Posterior Development. Current Biology, 19(5), R215–R219. 10.1016/j.cub.2009.01.052

Martinez-Barbera, J. P., Rodriguez, T. A., & Beddington, R. S. P. (2000). The Homeobox Gene Hesx1 Is Required in the Anterior Neural Ectoderm for Normal Forebrain Formation. Developmental Biology, 223(2), 422–430. 10.1006/DBIO.2000.9757

Mayran, A., Kolly, D., Lopez-Delisle, L., Romaniuk, Y., Leonardi, M., Cossy, A.-C., Lacroix, T., Rekaik, H., Amândio, A. R., Osteil, P., & Duboule, D. (2025). Cadherins modulate the self-organizing potential of pseudo-embryos. Cell Reports, 44(11), 116567. 10.1016/j.celrep.2025.116567

McNamara, H. M., Solley, S. C., Adamson, B., Chan, M. M., & Toettcher, J. E. (2024). Recording morphogen signals reveals mechanisms underlying gastruloid symmetry breaking. Nature Cell Biology 2024 26:11, 26(11), 1832–1844. 10.1038/s41556-024-01521-9

Moore-Scott, B. A., & Manley, N. R. (2005). Differential expression of Sonic hedgehog along the anterior-posterior axis regulates patterning of pharyngeal pouch endoderm and pharyngeal endoderm-derived organs. Developmental Biology, 278(2), 323–335. 10.1016/J.YDBIO.2004.10.027

Nandkishore, N., Vyas, B., Javali, A., Ghosh, S., & Sambasivan, R. (2018). Divergent early mesoderm specification underlies distinct head and trunk muscle programmes in vertebrates. Development (Cambridge), 145(18), 173187. 10.1242/DEV.160945/48477

Neijts, R., Amin, S., van Rooijen, C., & Deschamps, J. (2017). Cdx is crucial for the timing mechanism driving colinear Hox activation and defines a trunk segment in the Hox cluster topology. Developmental Biology, 422(2), 146–154. 10.1016/J.YDBIO.2016.12.024

Neijts, R., Simmini, S., Giuliani, F., van Rooijen, C., & Deschamps, J. (2014). Region-specific regulation of posterior axial elongation during vertebrate embryogenesis. Developmental Dynamics, 243(1), 88–98. 10.1002/DVDY.24027;ISSUE:ISSUE:DOI

Niederreither, K., Subbarayan, V., Dollé, P., & Chambon, P. (1999). Embryonic retinoic acid synthesis is essential for early mouse post-implantation development. Nature Genetics 1999 21:4, 21(4), 444–448. 10.1038/7788

Niehrs, C., Kazanskaya, O., Wu, W., & Glinka, A. (2001). Dickkopf1 and the Spemann-Mangold head organizer. In Int. J. Dev. Biol (Vol. 45). www.ijdb.ehu.es

Nolte, C., De Kumar, B., & Krumlauf, R. (2019). Hox genes: Downstream “effectors” of retinoic acid signaling in vertebrate embryogenesis. Genesis, 57(7), e23306. 10.1002/DVG.23306;ISSUE:ISSUE:DOI

Nowotschin, S., Ferrer-Vaquer, A., Concepcion, D., Papaioannou, V. E., & Hadjantonakis, A. K. (2012). Interaction of Wnt3a, Msgn1 and Tbx6 in neural versus paraxial mesoderm lineage commitment and paraxial mesoderm differentiation in the mouse embryo. Developmental Biology, 367(1), 1–14. 10.1016/J.YDBIO.2012.04.012

Pijuan-Sala, B., Griffiths, J. A., Guibentif, C., Hiscock, T. W., Jawaid, W., Calero-Nieto, F. J., Mulas, C., Ibarra-Soria, X., Tyser, R. C. V., Ho, D. L. L., Reik, W., Srinivas, S., Simons, B. D., Nichols, J., Marioni, J. C., & Göttgens, B. (2019). A single-cell molecular map of mouse gastrulation and early organogenesis. Nature 2019 566:7745, 566(7745), 490–495. 10.1038/s41586-019-0933-9

Pilon, N., Oh, K., Sylvestre, J. R., Bouchard, N., Savory, J., & Lohnes, D. (2006). Cdx4 is a direct target of the canonical Wnt pathway. Developmental Biology, 289(1), 55–63. 10.1016/J.YDBIO.2005.10.005

Pilon, N., Oh, K., Sylvestre, J. R., Savory, J. G. A., & Lohnes, D. (2007). Wnt signaling is a key mediator of Cdx1 expression in vivo. *Development (Cambridge*, England), 134(12), 2315–2323. 10.1242/DEV.001206

Ramkumar, N., & Anderson, K. V. (2011). SnapShot: Mouse Primitive Streak. Cell, 146(3), 488–488.e2. 10.1016/J.CELL.2011.07.028

Rawat, V. P. S., Thoene, S., Naidu, V. M., Arseni, N., Heilmeier, B., Metzeler, K., Petropoulos, K., Deshpande, A., Quintanilla-Martinez, L., Bohlander, S. K., Spiekermann, K., Hiddemann, W., Feuring-Buske, M., & Buske, C. (2008). Overexpression of CDX2 perturbs HOX gene expression in murine progenitors depending on its N-terminal domain and is closely correlated with deregulated HOX gene expression in human acute myeloid leukemia. Blood, 111(1), 309–319. 10.1182/BLOOD-2007-04-085407

Rayon, T., Menchero, S., Nieto, A., Xenopoulos, P., Crespo, M., Cockburn, K., Cañon, S., Sasaki, H., Hadjantonakis, A. K., de la Pompa, J. L., Rossant, J., & Manzanares, M. (2014). Notch and Hippo Converge on Cdx2 to Specify the Trophectoderm Lineage in the Mouse Blastocyst. Developmental Cell, 30(4), 410–422. 10.1016/j.devcel.2014.06.019

Rekaik, H., Lopez-Delisle, L., Hintermann, A., Mascrez, B., Bochaton, C., Mayran, A., & Duboule, D. (2023). Sequential and directional insulation by conserved CTCF sites underlies the Hox timer in stembryos. Nature Genetics 2023 55:7, 55(7), 1164–1175. 10.1038/s41588-023-01426-7

Rhinn, M., & Dollé, P. (2012). Retinoic acid signalling during development. *Development (Cambridge*, England), 139(5), 843–858. 10.1242/DEV.065938

Robertson, E. J. (2014). Dose-dependent Nodal/Smad signals pattern the early mouse embryo. Seminars in Cell & Developmental Biology, 32, 73–79. 10.1016/J.SEMCDB.2014.03.028

Rosen, L. U., Stapel, L. C., Argelaguet, R., Barker, C. G., Yang, A., Reik, W., & Marioni, J. C. (2022). Inter-gastruloid heterogeneity revealed by single cell transcriptomics time course: implications for organoid based perturbation studies. BioRxiv, 2022.09.27.509783. 10.1101/2022.09.27.509783

Rossant, J., Zirngibl, R., Cado, D., Shago, M., & Giguère, V. (1991). Expression of a retinoic acid response element-hsplacZ transgene defines specific domains of transcriptional activity during mouse embryogenesis. Genes & Development, 5(8), 1333–1344. 10.1101/GAD.5.8.1333

Rossi, G., Broguiere, N., Miyamoto, M., Boni, A., Guiet, R., Girgin, M., Kelly, R. G., Kwon, C., & Lutolf, M. P. (2021). Capturing Cardiogenesis in Gastruloids. Cell Stem Cell, 28(2), 230–240.e6. 10.1016/j.stem.2020.10.013

Ruiz I Altaba, A., & Jessell, T. (1991). Retinoic acid modifies mesodermal patterning in early Xenopus embryos. Genes & Development, 5(2), 175–187. 10.1101/GAD.5.2.175

Ruiz I Altaba, S., & Jessell, T. M. (1991). Retinoic acid modifies the pattern of cell differentiation in the central nervous system of neurula stage Xenopus embryos. Development, 112(4), 945–958. 10.1242/DEV.112.4.945

Russ, A. P., Wattler, S., Colledge, W. H., Aparicio, S. A. J. R., Carlton, M. B. L., Pearce, J. J., Barton, S. C., Azim Surani, M., Ryan, K., Nehls, M. C., Wilsons, V., & Evans, M. J. (2000). Eomesodermin is required for mouse trophoblast development and mesoderm formation. Nature 2000 404:6773, 404(6773), 95–99. 10.1038/35003601

Sambasivan, R., Gayraud-Morel, B., Dumas, G., Cimper, C., Paisant, S., Kelly, R., & Tajbakhsh, S. (2009). Distinct Regulatory Cascades Govern Extraocular and Pharyngeal Arch Muscle Progenitor Cell Fates. Developmental Cell, 16(6), 810–821. 10.1016/j.devcel.2009.05.008

Sambasivan, R., & Steventon, B. (2021). Neuromesodermal Progenitors: A Basis for Robust Axial Patterning in Development and Evolution. Frontiers in Cell and Developmental Biology, 8, 607516. 10.3389/FCELL.2020.607516/BIBTEX

Schubert, M., Holland, N. D., Laudet, V., & Holland, L. Z. (2006). A retinoic acid-Hox hierarchy controls both anterior/posterior patterning and neuronal specification in the developing central nervous system of the cephalochordate amphioxus. Developmental Biology, 296(1), 190–202. 10.1016/J.YDBIO.2006.04.457

Schüle, K. M., Weckerle, J., Probst, S., Wehmeyer, A. E., Zissel, L., Schröder, C. M., Tekman, M., Kim, G. J., Schlägl, I. M., Sagar, & Arnold, S. J. (2023). Eomes restricts Brachyury functions at the onset of mouse gastrulation. Developmental Cell, 58(18), 1627–1642.e7. 10.1016/j.devcel.2023.07.023

Schwaiger, M., Andrikou, C., Dnyansagar, R., Murguia, P. F., Paganos, P., Voronov, D., Zimmermann, B., Lebedeva, T., Schmidt, H. A., Genikhovich, G., Benvenuto, G., Arnone, M. I., & Technau, U. (2022). An ancestral Wnt-Brachyury feedback loop in axial patterning and recruitment of mesoderm-determining target genes. Nature Ecology & Evolution, 6(12), 1921–1939. 10.1038/S41559-022-01905-W

Serup, P., Gustavsen, C., Klein, T., Potter, L. A., Lin, R., Mullapudi, N., Wandzioch, E., Hines, A., Davis, A., Bruun, C., Engberg, N., Petersen, D. R., Peterslund, J. M. L., MacDonald, R. J., Grapin-Botton, A., Magnuson, M. A., & Zaret, K. S. (2012). Partial promoter substitutions generating transcriptional sentinels of diverse signaling pathways in embryonic stem cells and mice. DMM Disease Models and Mechanisms, 5(6), 956–966. 10.1242/DMM.009696/258717/AM/PARTIAL-PROMOTER-SUBSTITUTIONS-GENERATING

Simeone, A., & Acampora, D. (2001). Otx2 function in murine gastrulation The role of Otx2 in organizing the anterior patterning in mouse. In Int. J. Dev. Biol (Vol. 45). www.ijdb.ehu.es

Simon, C. S., Downes, D. J., Gosden, M. E., Telenius, J., Higgs, D. R., Hughes, J. R., Costello, I., Bikoff, E. K., & Robertson, E. J. (2017). Functional characterisation of cis-regulatory elements governing dynamic Eomes expression in the early mouse embryo. Development (Cambridge), 144(7), 1249–1260. 10.1242/DEV.147322/264330/AM/FUNCTIONAL-CHARACTERISATION-OF-CIS-REGULATORY

Smith, J. C., Price, B. M. J., Green, J. B. A., Weigel, D., & Herrmann, B. G. (1991). Expression of a xenopus homolog of Brachyury (T) is an immediate-early response to mesoderm induction. Cell, 67(1), 79–87. 10.1016/0092-8674(91)90573-H

Stern, C. D., Charité, J., Deschamps, J., Duboule, D., Durston, A. J., Kmita, M., Nicolas, J.-F., Palmeirim, I., Smith, J. C., & Wolpert, L. (2006). Head-tail patterning of the vertebrate embryo: one, two or many unresolved problems? The International Journal of Developmental Biology, 50(1), 3–15. 10.1387/ijdb.052095cs

Suppinger, S., Zinner, M., Aizarani, N., Lukonin, I., Ortiz, R., Azzi, C., Stadler, M. B., Vianello, S., Palla, G., Kohler, H., Mayran, A., Lutolf, M. P., & Liberali, P. (2023). Multimodal characterization of murine gastruloid development. Cell Stem Cell, 30(6), 867–884.e11. 10.1016/J.STEM.2023.04.018

Sussel, L., Marin, O., Kimura, S., & Rubenstein, J. L. R. (1999). Loss of Nkx2.1 homeobox gene function results in a ventral to dorsal molecular respecification within the basal telencephalon: evidence for a transformation of the pallidum into the striatum. Development, 126(15), 3359–3370. 10.1242/DEV.126.15.3359

Takada, S., Stark, K. L., Shea, M. J., Vassileva, G., McMahon, J. A., & McMahon, A. P. (1994). Wnt-3a regulates somite and tailbud formation in the mouse embryo. Genes & Development, 8(2), 174–189. 10.1101/GAD.8.2.174

Takemoto, T., Uchikawa, M., Yoshida, M., Bell, D. M., Lovell-Badge, R., Papaioannou, V. E., & Kondoh, H. (2011). Tbx6-dependent Sox2 regulation determines neural or mesodermal fate in axial stem cells. Nature 2011 470:7334, 470(7334), 394–398. 10.1038/nature09729

Thomas, P., & Beddington, R. (1996). Anterior primitive endoderm may be responsible for patterning the anterior neural plate in the mouse embryo. Current Biology, 6(11), 1487–1496. 10.1016/S0960-9822(96)00753-1

Thomas, P. Q., Dattani, M. T., Brickman, J. M., McNay, D., Warne, G., Zacharin, M., Cameron, F., Hurst, J., Woods, K., Dunger, D., Stanhope, R., Forrest, S., Robinson, I. C. A. F., & Beddington, R. S. P. (2001). Heterozygous HESX1 mutations associated with isolated congenital pituitary hypoplasia and septo-optic dysplasia. Human Molecular Genetics, 10(1), 39–45. 10.1093/HMG/10.1.39

Tian, T., & Meng, A. M. (2006). Nodal signals pattern vertebrate embryos. Cellular and Molecular Life Sciences, 63(6), 672–685. 10.1007/S00018-005-5503-7/METRICS

Tirosh, I., Izar, B., Prakadan, S. M., Wadsworth, M. H., Treacy, D., Trombetta, J. J., Rotem, A., Rodman, C., Lian, C., Murphy, G., Fallahi-Sichani, M., Dutton-Regester, K., Lin, J. R., Cohen, O., Shah, P., Lu, D., Genshaft, A. S., Hughes, T. K., Ziegler, C. G. K.,…Garraway, L. A. (2016). Dissecting the multicellular ecosystem of metastatic melanoma by single-cell RNA-seq. Science, 352(6282), 189–196. 10.1126/SCIENCE.AAD0501;JOURNAL:JOURNAL:SCIENCE;WGROUP:ST RING:PUBLICATION

Traag, V. A., Waltman, L., & van Eck, N. J. (2019). From Louvain to Leiden: guaranteeing well-connected communities. Scientific Reports, 9(1). 10.1038/S41598-019-41695-Z

Turner, D. A., Girgin, M., Alonso-Crisostomo, L., Trivedi, V., Baillie-Johnson, P., Glodowski, C. R., Hayward, P. C., Collignon, J., Gustavsen, C., Serup, P., Steventon, B., Lutolf, M. P., & Arias, A. M. (2017). Anteroposterior polarity and elongation in the absence of extraembryonic tissues and of spatially localised signalling in gastruloids: Mammalian embryonic organoids. Development (Cambridge), 144(21), 3894–3906. 10.1242/DEV.150391/VIDEO-2

Turner, D. A., Rué, P., Mackenzie, J. P., Davies, E., & Martinez Arias, A. (2014). Brachyury cooperates with Wnt/β-catenin signalling to elicit primitive-streak-like behaviour in differentiating mouse embryonic stem cells. BMC Biology 2014 12:1, 12(1), 63-. 10.1186/S12915-014-0063-7

Uehara, M., Yashiro, K., Takaoka, K., Yamamoto, M., & Hamada, H. (2009). Removal of maternal retinoic acid by embryonic CYP26 is required for correct Nodal expression during early embryonic patterning. Genes & Development, 23(14), 1689–1698. 10.1101/GAD.1776209

van de Ven, C., Bialecka, M., Neijts, R., Young, T., Rowland, J. E., Stringer, E. J., van Rooijen, C., Meijlink, F., Nóvoa, A., Freund, J. N., Mallo, M., Beck, F., & Deschamps, J. (2011). Concerted involvement of Cdx/Hox genes and Wnt signaling in morphogenesis of the caudal neural tube and cloacal derivatives from the posterior growth zone. Development, 138(16), 3451–3462. 10.1242/DEV.066118

van den Akker, E., Forlani, S., Chawengsaksophak, K., de Graaft, W., Beck, F., Meyer, B. I., & Deschamps, J. (2002). Cdx1 and Cdx2 have overlapping functions in anteroposterior patterning and posterior axis elongation. Development, 129(9), 2181–2193. 10.1242/DEV.129.9.2181

Van Den Ameele, J., Tiberi, L., Bondue, A., Paulissen, C., Herpoel, A., Iacovino, M., Kyba, M., Blanpain, C., & Vanderhaeghen, P. (2012). Eomesodermin induces Mesp1 expression and cardiac differentiation from embryonic stem cells in the absence of Activin. EMBO Reports, 13(4), 355–362. 10.1038/EMBOR.2012.23/FIGURES/4

van den Brink, S. C., Alemany, A., van Batenburg, V., Moris, N., Blotenburg, M., Vivié, J., Baillie-Johnson, P., Nichols, J., Sonnen, K. F., Martinez Arias, A., & van Oudenaarden, A. (2020). Single-cell and spatial transcriptomics reveal somitogenesis in gastruloids. Nature 2020 582:7812, 582(7812), 405–409. 10.1038/s41586-020-2024-3

van Rooijen, C., Simmini, S., Bialecka, M., Neijts, R., van de Ven, C., Beck, F., & Deschamps, J. (2012). Evolutionarily conserved requirement of Cdx for post-occipital tissue emergence. Development, 139(14), 2576–2583. 10.1242/DEV.079848

Vyas, B., Nandkishore, N., & Sambasivan, R. (2019). Vertebrate cranial mesoderm: developmental trajectory and evolutionary origin. Cellular and Molecular Life Sciences 2019 77:10, 77(10), 1933–1945. 10.1007/S00018-019-03373-1

Wang, W. C. H., & Shashikant, C. S. (2007). Evidence for positive and negative regulation of the mouse Cdx2 gene. Journal of Experimental Zoology Part B: Molecular and Developmental Evolution, 308B(3), 308–321. 10.1002/JEZ.B.21154

Wehmeyer, A. E., Schmitt, J. K., Eggersdorfer, F., Zissel, L., Schröder, C. M., Tekman, M., Dias, A., Schüle, K. M., Probst, S., Martinez-Arias, A., McDole, K., & Arnold, S. J. (2025). Competing regulatory modules control the transition between mammalian gastrulation modes. BioRxiv, 2025.05.07.652670. 10.1101/2025.05.07.652670

Wolf, F. A., Angerer, P., & Theis, F. J. (2018). SCANPY: large-scale single-cell gene expression data analysis. Genome Biology 2018 19:1, 19(1), 15-. 10.1186/S13059-017-1382-0

Wolock, S. L., Lopez, R., & Klein, A. M. (2019). Scrublet: Computational Identification of Cell Doublets in Single-Cell Transcriptomic Data. Cell Systems, 8(4), 281–291.e9. 10.1016/j.cels.2018.11.005

Wu, C., MacLeod, I., & Su, A. I. (2013). BioGPS and MyGene.info: organizing online, gene-centric information. Nucleic Acids Research, 41(D1), D561–D565. 10.1093/NAR/GKS1114

Yamaguchi, T. P. (2001). Heads or tails: Wnts and anterior-posterior patterning. Current Biology, 11(17), R713–R724. 10.1016/S0960-9822(01)00417-1

Yamaguchi, T. P., Yoshikawa, Y., Wu, N., Mcmahon, A. P., & Takada, S. (1999). T (Brachyury) is a direct target of Wnt3a during paraxial mesoderm specification. Genes &Development, 13(24), 3185–3190. 10.1101/gad.13.24.3185

Young, T., & Deschamps, J. (2009). Chapter 8 Hox, Cdx, and Anteroposterior Patterning in the Mouse Embryo. Current Topics in Developmental Biology, 88, 235–255. 10.1016/S0070-2153(09)88008-3

Young, T., Rowland, J. E., van de Ven, C., Bialecka, M., Novoa, A., Carapuco, M., van Nes, J., de Graaff, W., Duluc, I., Freund, J. N., Beck, F., Mallo, M., & Deschamps, J. (2009). Cdx and Hox Genes Differentially Regulate Posterior Axial Growth in Mammalian Embryos. Developmental Cell, 17(4), 516–526. 10.1016/j.devcel.2009.08.010

Zou, D., Silvius, D., Davenport, J., Grifone, R., Maire, P., & Xu, P. X. (2006). Patterning of the third pharyngeal pouch into thymus/parathyroid by Six and Eya1. Developmental Biology, 293(2), 499–512. 10.1016/J.YDBIO.2005.12.015

