## Supplementary figures and methods for "Retinoic acid coordinates the orderly construction of the mammalian body in the anterior-to-posterior sequence" (from another version of this preprint: version 1, 2025-12-13)

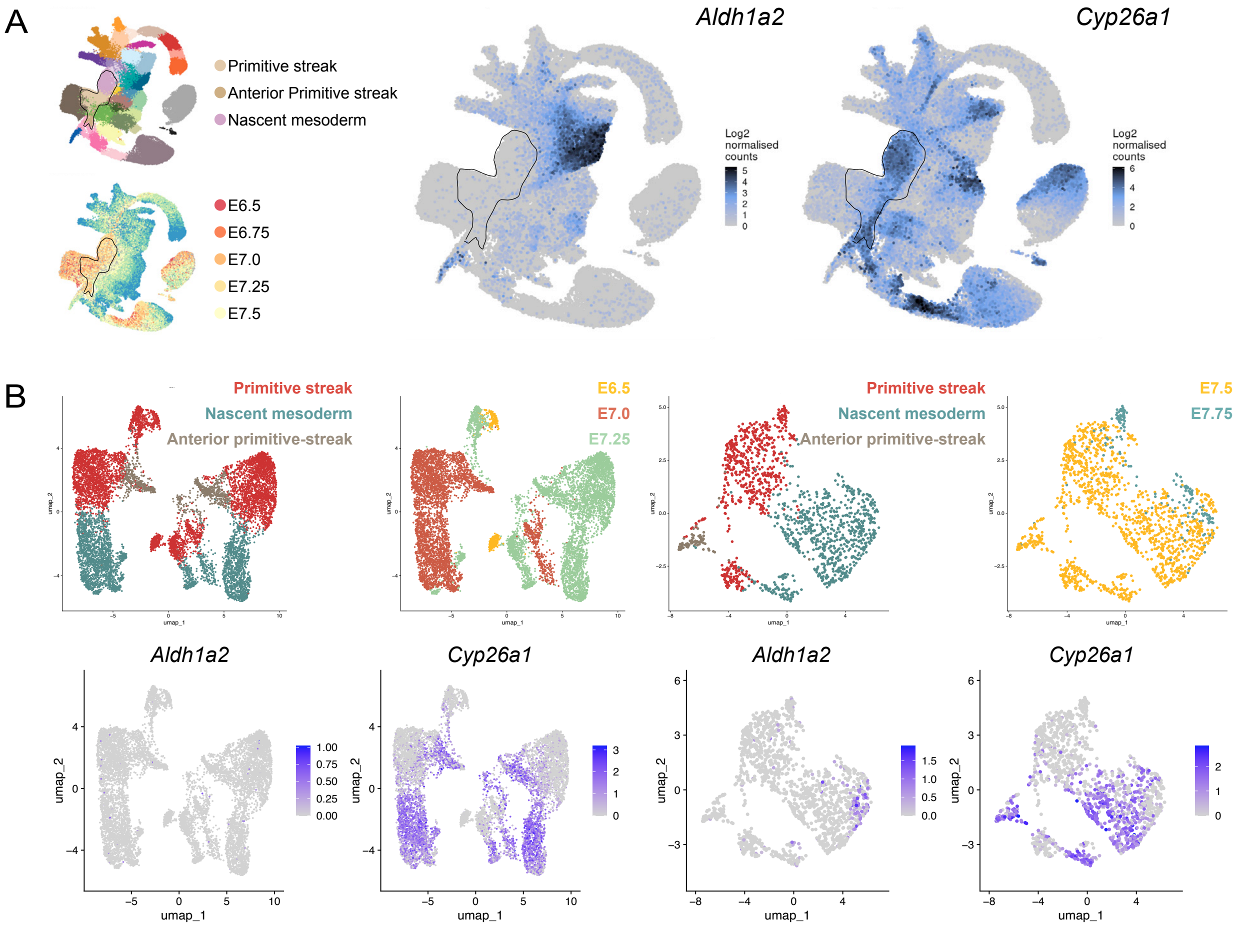
A
Primitive streak
Anterior Primitive streak
Nascent mesoderm
E6.5
E6.75
E7.0
E7.25
E7.5
Aldh1a2
Cyp26a1
Log2 normalised counts
B
Primitive streak
Nascent mesoderm
Anterior primitive-streak
E6.5
E7.0
E7.25
E7.5
E7.75
Aldh1a2
Cyp26a1
umap_1
umap_2

**Figure S1: RA signaling is excluded during anterior mesoderm specification**

(A) Analysis of the mouse gastrulation ATLAS (Pijuan-Sala et al, 2019). The dataset overview is reproduced from https://crukci.shinyapps.io/mousegastrulation2018/ for reference. The primitive streak, anterior primitive streak, nascent mesoderm, and mixed mesoderm clusters, as annotated in their article, are marked with a black boundary line. Expression data for *Aldh1a2* and *Cyp26a1* are projected on the UMAP.

(B) Subset of primitive streak, anterior primitive streak, nascent mesoderm, and mixed mesoderm clusters from E6.5 to E7.25 (left) and E7.5 to E7.75 (right) showing expression of *Aldh1a2* and *Cyp26a1*.

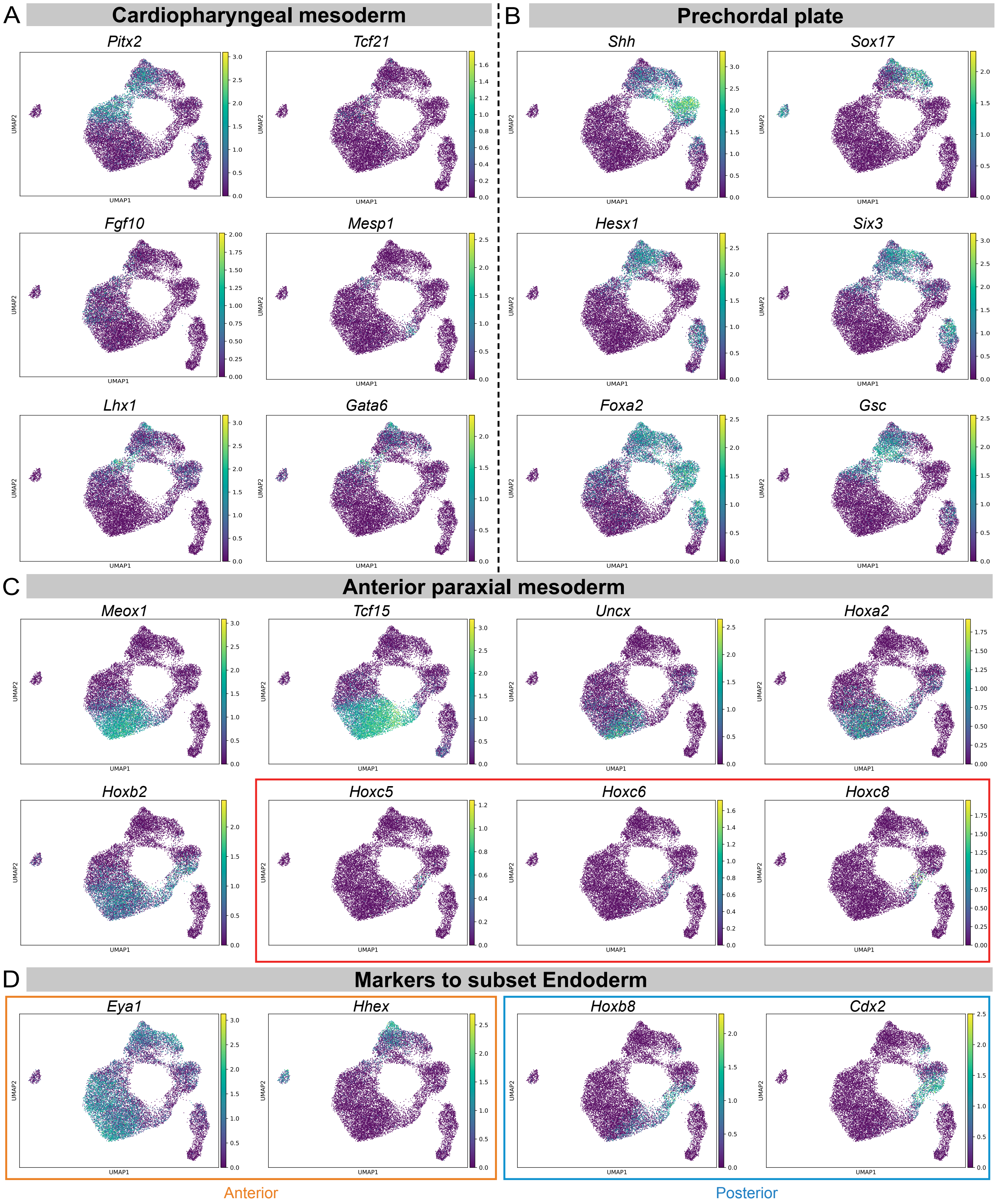

A
Cardiopharyngeal mesoderm
Pitx2
Tcf21
Fgf10
Mesp1
Lhx1
Gata6
B
Prechordal plate
Shh
Sox17
Hesx1
Six3
Foxa2
Gsc
C
Anterior paraxial mesoderm
Meox1
Tcf15
Uncx
Hoxa2
Hoxb2
Hoxc5
Hoxc6
Hoxc8
D
Markers to subset Endoderm
Eya1
Hhex
Hoxb8
Cdx2
Anterior
Posterior
UMAP1
UMAP2

**Figure S2: Marker expression domains reveal cluster identities in wild-type single-cell transcriptomic data**

(A) Gene interrogation plots: Expression of *Pitx2, Tcf21, Fgf10, Mesp1, Lhx1,* and *Gata6* marks the cardiopharyngeal mesoderm in wild type sc-RNA seq UMAP shown in Figure 2.

(B) Gene interrogation plots: Expression of *Shh, Sox17, Hesx1, Six3, Foxa2,* and *Gsc* marks the prechordal plate cluster.

(C) Gene interrogation plots: The presence of *Meox1, Tcf15, Uncx, Hoxa2,* and *Hoxb2,* and the absence (indicated by a red box) of *Hoxc5, Hoxc6,* and *Hoxc8* expression, mark the anterior paraxial mesoderm cluster.

(D) Gene interrogation plots: Expression of *Eya1, Hhex* marks the anterior endodermal subcluster (marked in orange box), and *Hoxb8, Cdx2* mark the posterior endodermal cluster (marked in blue box).

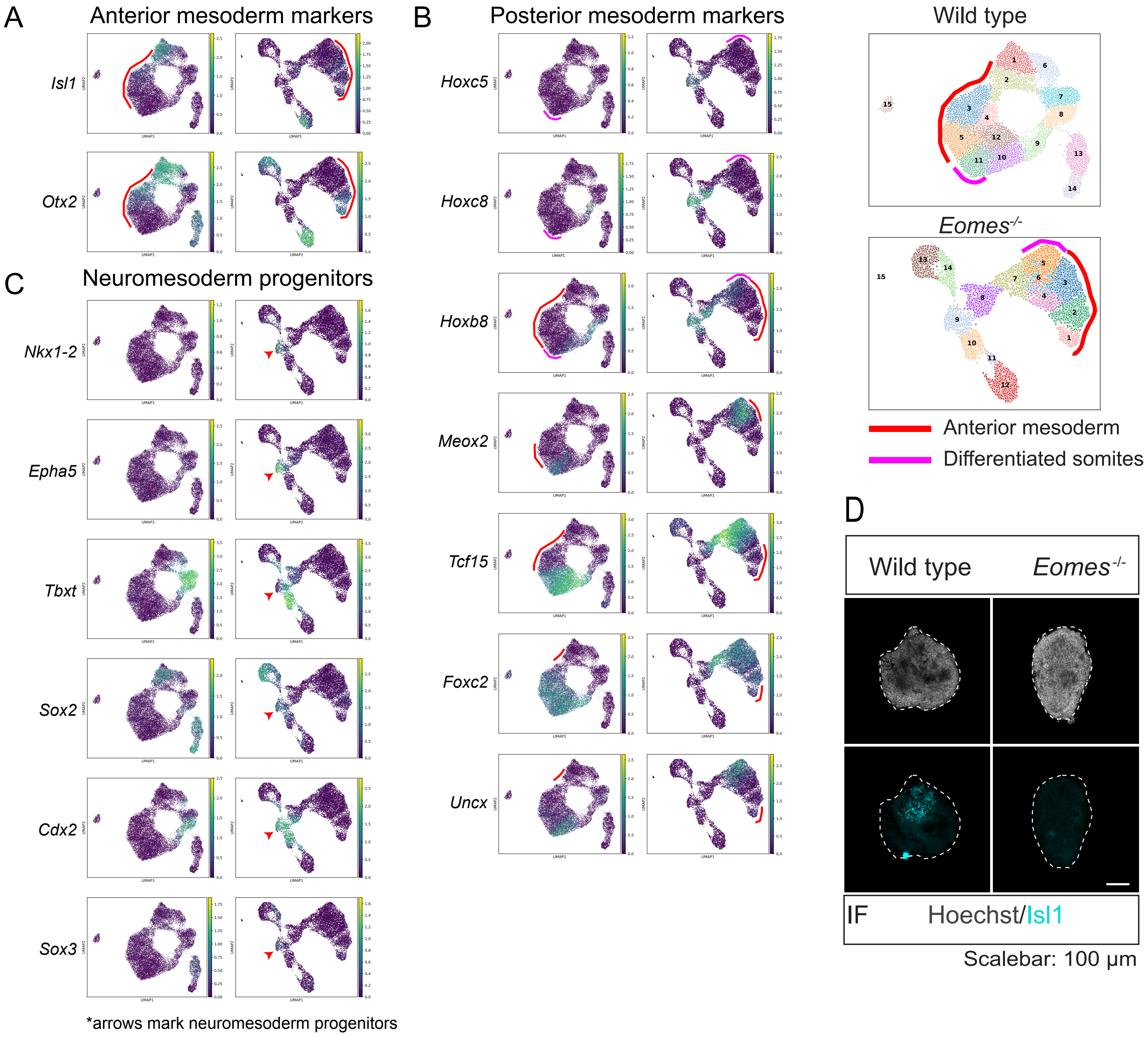

A
Anterior mesoderm markers
Isl1
Otx2
B
Posterior mesoderm markers
Hoxc5
Hoxc8
Hoxb8
Meox2
Tcf15
Foxc2
Uncx
Wild type
Eomes-/-
Anterior mesoderm
Differentiated somites
C
Neuromesoderm progenitors
Nkx1-2
Epha5
Tbxt
Sox2
Cdx2
Sox3
*arrows mark neuromesoderm progenitors
D
Wild type
Eomes-/-
IF
Hoechst/Isl1
Scalebar: 100 µm

**Figure S3: Marker expression domains reveal cluster identities in *Eomes*$^{-/-}$ single-cell transcriptomic data**

(A) Gene interrogation plots: Expression of *Isl1 and Otx2,* which mark the cardiopharyngeal mesoderm and anterior mesoderm, is expressed in low levels in the anterior mesoderm clusters (boundary marked in red for corresponding clusters, both in wild type and *Eomes*$^{-/-}$) in *Eomes*$^{-/-}$ gastruloids, compared with their expression in wild type.

(B) Gene interrogation plots: Expression of *Nkx1-2, Epha5, Tbxt, Sox2, Cdx2, and Sox3,* which mark the neuromesoderm progenitors (NMP). The NMP cluster in *Eomes*$^{-/-}$ plots is marked with an arrow.

(C) Gene interrogation plots: Expression of posterior mesoderm markers *Hoxc5, Hoxc8, Hoxb8, Meox2, Tcf15, Foxc2, and Uncx.* The boundary for the corresponding clusters in wild type and mutant is marked with red for the anterior mesoderm and pink for differentiated somites.

The annotations for the marked clusters are as follows. Wild type: 2-Cardiopharyngeal mesoderm, 3-Pharyngeal mesoderm 1, 4-Pharyngeal mesoderm 2, 5-Anterior paraxial mesoderm, 11-Dermomyotome, 12-Sclerotome; *Eomes*$^{-/-}$: 1-Cardiac mesoderm, 2-Pharyngeal mesoderm, 3-Anterior paraxial mesoderm 1, 4-Anterior paraxial mesoderm 2, 5-Dermomyotome, 6-Sclerotome

(D) Confocal images of gastruloids immunostained for the marker Isl1. Wild type AM gastruloids (N = 3, n = 11), *Eomes*$^{-/-}$ AM (N = 3, n = 14).

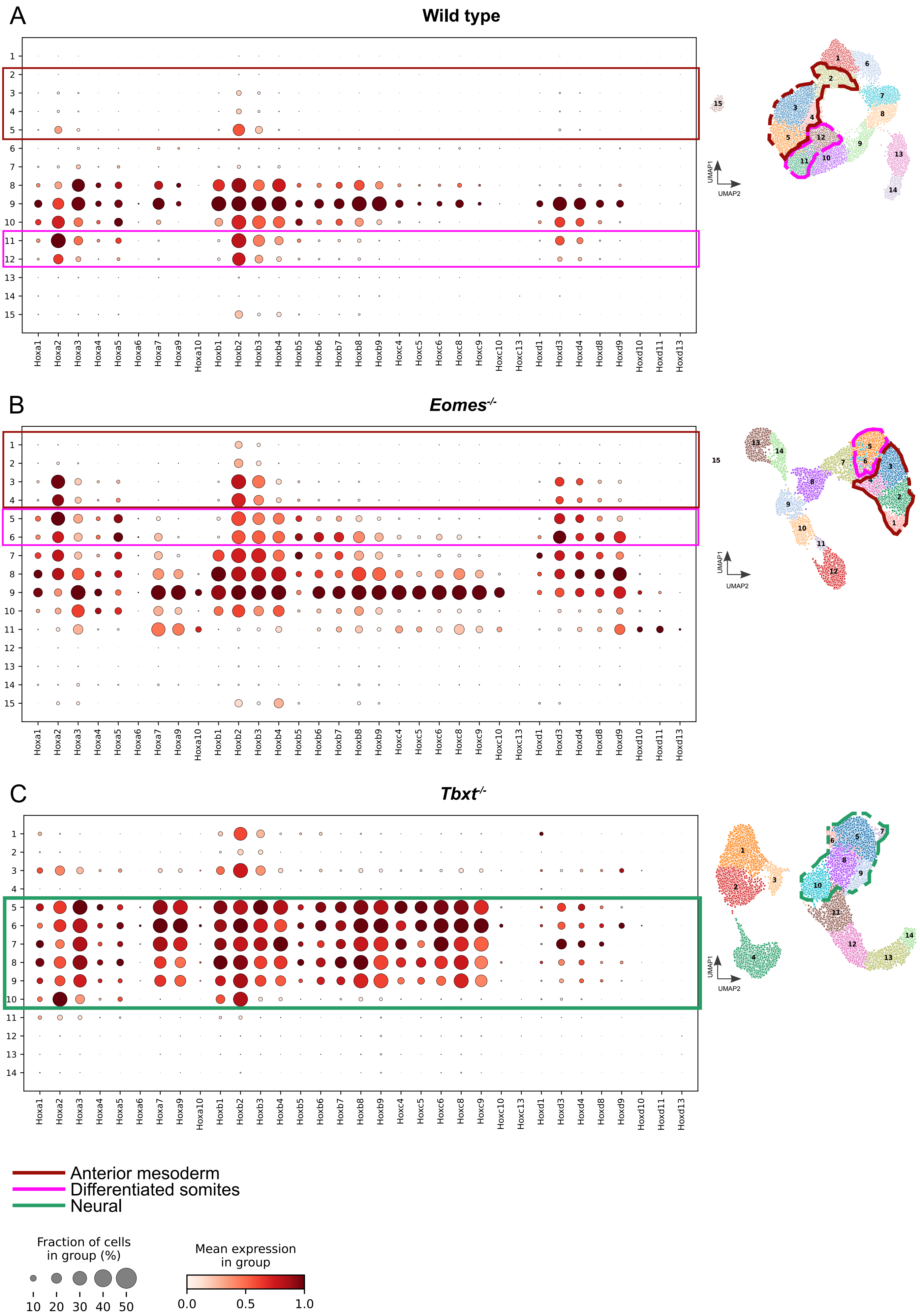

A
Wild type
B
Eomes-/-
C
Tbxt-/-
Hoxa1 Hoxa2 Hoxa3 Hoxa4 Hoxa5 Hoxa6 Hoxa7 Hoxa9 Hoxa10 Hoxb1 Hoxb2 Hoxb3 Hoxb4 Hoxb5 Hoxb6 Hoxb7 Hoxb8 Hoxb9 Hoxc4 Hoxc5 Hoxc6 Hoxc8 Hoxc9 Hoxc10 Hoxc13 Hoxd1 Hoxd3 Hoxd4 Hoxd8 Hoxd9 Hoxd10 Hoxd11 Hoxd13
UMAP1
UMAP2
Anterior mesoderm
Differentiated somites
Neural
Fraction of cells in group (%)
10 20 30 40 50
Mean expression in group
0.0 0.5 1.0

**Figure S4: *Eomes* and *Tbxt* mutants acquire posterior identity**

(A) Dot plot of *Hox* gene expression in wild type single-cell transcriptome data set.

(B) Dot plot of *Hox* gene expression in *Eomes*$^{-/-}$ single-cell transcriptome.

(C) Dot plot of *Hox* gene expression in *Tbxt*$^{-/-}$ single-cell transcriptome.

The anterior mesoderm clusters are highlighted by a red rectangle, and the differentiated somite clusters are highlighted in pink. The neural clusters are highlighted by a green rectangle.

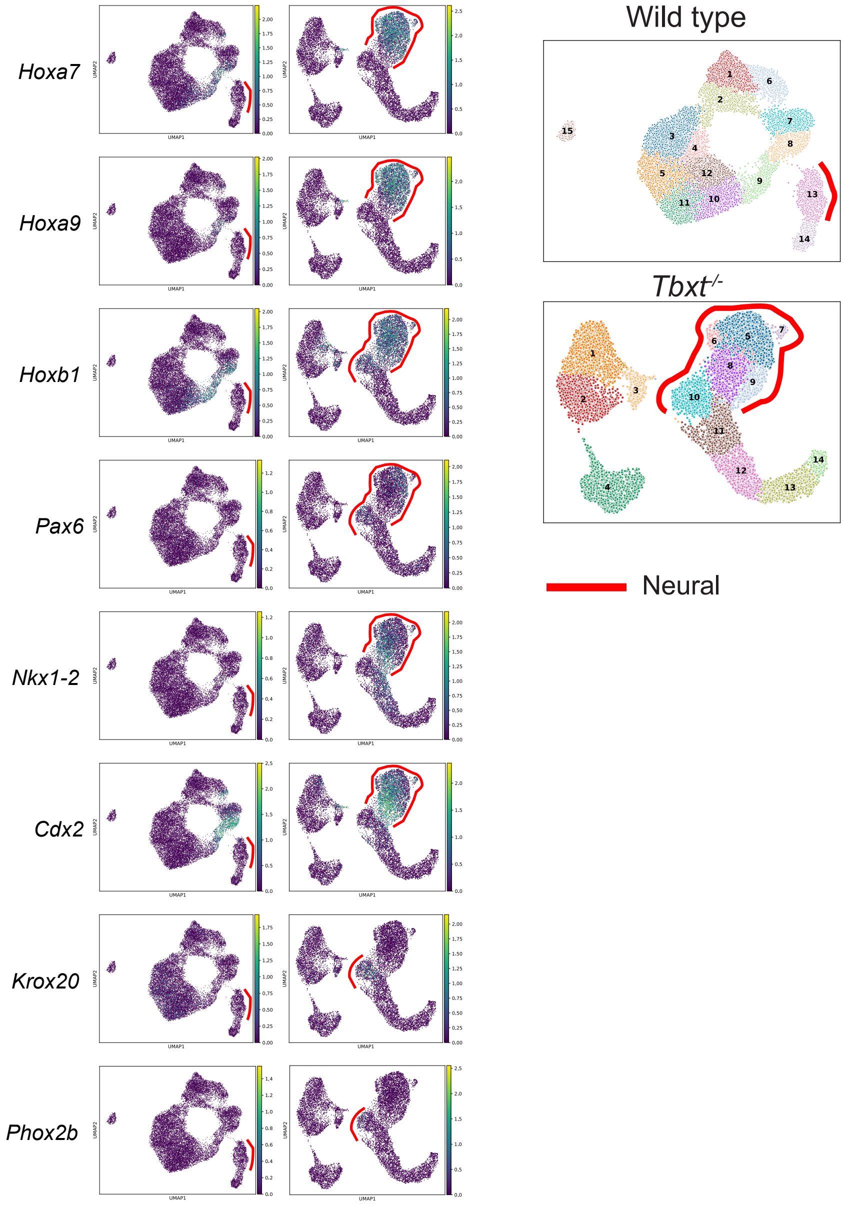
Hoxa7
Hoxa9
Hoxb1
Pax6
Nkx1-2
Cdx2
Krox20
Phox2b
Wild type
Tbxt-/-
Neural

**Figure S5: Absence of *Tbxt* leads to the formation of posterior neural populations**

Gene interrogation plots: *Hoxa7, Hoxa9, and Hoxb1* are associated with neural development. *Pax6, Nkx1-2, and Cdx2* mark posterior neural development. *Krox20* and *Phox2b* are hindbrain markers. The boundary for the cluster is marked with a red line, highlighting neural populations.

Annotations of the highlighted clusters are as follows. Wild type: 13-Anterior neurectoderm; *Tbxt*$^{-/-}$: 5-Spinal cord, 6-Dorsal neural tube, 7-Neural progenitors, 8-Neuromesoderm progenitors, 9-Neurally-committed neuromesoderm progenitors, 10-Hindbrain

### SUPPLEMENTARY METHODS

#### Generation and validation of *RARE-d2-eGFP* reporter line

The pGL3 RARE-d2EGFP plasmid (Addgene #183055; gift from Chaya Kalcheim) was used to generate a retinoic acid-responsive mESC reporter line. mESCs were seeded at 1 × $10^5$ cells/mL on a 0.1% gelatin-coated 24-well plate and transfected with 1µg plasmid DNA, 0.75µL Lipofectamine 3000, and 2µL P3000 reagent (L3000-008, Invitrogen), each diluted in 25µL serum-free medium. The mixtures were combined, incubated for 15 min at room temperature, and added dropwise to the cells. After 24h, the medium was refreshed, and the cells were cultured for another 24h before passaging.

Cells were then seeded at clonal density (0.5 × $10^5$ cells/mL) in 0.1% gelatin-coated 60-mm dishes and cultured for ~10 days until individual colonies formed. Colonies were manually picked, dissociated with TrypLE (5 min, 37°C), and expanded in 48-well plates. Clones were screened by PCR using primers. RARE-d2-eGFP gastruloids display dispersed reporter activity.

F: 5′-GGGTAGGGTTCACCGAAAGT-3′ and

R: 5′-GACTGGGTGCTCAGGTAGTG-3′

#### Generation of E14Tg2a-GFP mESCs

Lentiviral GFP reporter particles were produced by cotransfection of pRRL-SIN-PGK-EGFP (Addgene #12252), psPAX2 (Addgene #12260), and pMD2.G (Addgene #12259) plasmids into HEK293T-LX packaging cells using a lipofectamine-based transfection. After 48h, virus-containing supernatants were collected and concentrated using Vivaspin columns (100kDa MWCO; VC1042, Sartorius) according to the manufacturer's instructions. Concentrated viral aliquots were snap-frozen in liquid nitrogen and stored at −80°C. Viral titer determined using the following formula: Titer (TU/mL) = (N × C × D)/0.005, where N is the average number of clones per field of view, C is the ratio of the well area to the field of view area, and D is the fold dilution of the viral vector stock. For transduction, 5 × $10^4$ E14 mESCs were plated on a 0.1% gelatin-coated 4-well dish. After 24h, the growth medium was replaced with fresh medium containing the desired viral dilution and 8µg/mL polybrene (TR-1003, Merck). Following a 24h incubation, the viral media was removed, and the cells were rinsed with PBS and then cultured in standard growth medium. Once confluent, cells were dissociated and reseeded at clonal density. Individual GFP-positive colonies were screened by epifluorescence microscopy (Olympus IX83), manually picked, expanded, and validated for pluripotency and gastruloid-forming potential prior to use in experiments.

#### Generation of *Eomes* and *Tbxt* null mutant mESC lines

Loss-of-function mutations in the T-box genes *Eomes* and *Tbxt* were generated using CRISPR–Cas9–mediated genome editing. sgRNAs were designed to target the start codon and the T-box DNA-binding domain using the CRISPR design tool (crispr.mit.edu). E14Tg2a mESCs were plated at a density of 2.6 × $10^4$ cells/cm² and transfected at ~50% confluency with two pX459 plasmids, each containing an sgRNA and a Cas9, puromycin-resistance cassette, using Lipofectamine 3000 (Gibco; Cat. L3000008). After 24h, the transfection mix was removed, and cells were allowed to recover for an additional 24h. Cells were trypsinized and plated at low density (7 × $10^3$ cells/cm²). Puromycin selection (1µg/mL; Sigma, P8833) was applied for 4 days, after which surviving cells were expanded to obtain single-cell–derived colonies.

Colonies containing ~50–80 cells were mechanically picked under a microscope, dissociated with trypsin, and expanded in 48-well plates. Each clone was split into duplicate plates: one for genotyping and one for cryopreservation. For genotyping, genomic DNA was amplified using primer pairs flanking the expected deletion to detect the edited allele, and deletion-specific primers to confirm homozygosity. Clones carrying biallelic deletions were further validated by loss of protein expression using Western blotting and/or immunostaining for Eomes and Tbxt. The *Eomes* deletion spans chr9:118,307,912–118,308,601 (GRCm39), corresponding to bases 501–1190 of the genotyping PCR amplicon. The *Tbxt* deletion spans chr17:8,653,347–8,653,810 (GRCm39), corresponding to bases 86–549 of the genotyping PCR amplicon. Karyotyping was performed following Colcemid treatment (0.1µg/ mL, Invitrogen, Catalogue number: 15212012), and clones with a normal karyotype in at least 11 of 20 spreads (KOMP/UC Davis guidelines) were retained for downstream validation.

Potential off-target sites for each sgRNA were predicted using CHOPCHOP. For each predicted off-target locus, ~200 bp flanking the expected cut site were PCR-amplified from genomic DNA of the mutant clones and analyzed by Sanger sequencing. Perfect alignment with the mouse reference genome confirmed the absence of detectable off-target mutations. The predicted off targets and the results of the screen are listed in table S1.

**Methods table S1**

| Guide RNA | Target | Predicted off-target sequence | Predicted off-target location | Results |
|---|---|---|---|---|
| CGGCGCGCACTTCTACTCGC | *Eomes* | No predicted off-targets | - | - |
| TACGCCGGAACCTCAGCGGC | *Eomes* | CCTGaCGCTGAGGTTCaGGCtTA | chr2:165611264 | Not mutated |
| GAGTCAAGACTCCTGGAAGG | *Tbxt* | CCAgCTTCCAGGAGTCTTtACTg | chr1:114026062 | Not verified |
| | | GAGgCAAGACTCCTGGAgtGTGG | chr1:121060453 | Not mutated |

| | | | | |
|---|---|---|---|---|
| | | CCTCaTaCCAGG AaTCTTGACTC | chr11:102520906 | Not mutated |
| | | CCACCTTCCAGGt GTCTTGAacC | chr14:110065317 | Not verified |
| | | GAGTgAAaACTCC TGtAAGGGGG | chr16:79995669 | Not mutated |
| | | CCGCCTTCCAGa AGaCTTGcCTC | chr18:37752525 | Not verified |
| | | CCTCCTTCCAGG AGcCTTGgtTC | chr19:40657095 | Not mutated |
| | | CCACCTTtCAGGA GTCTcaACTC | chr2:62402934 | Not mutated |
| | | CCCCgTTCCAGG AGcCTgGACTC | chr2:163339721 | Not verified |
| | | GAcTCAgGACTCC TtGAAGGTGG | chr2:180470353 | Not mutated |
| | | GAGTCcAGgCTC CTGGAAGcTGG | chr4:119151311 | Not mutated |
| | | GgtTgAAGACTCC TGGAAGGTGG | chr6:133727944 | Not mutated |
| GAATGACCAGGTTTGCCTAG | *Tbxt* | GAATGACCAaGT TaGCCTAtAGG | chr1:32052621 | Not mutated |
| | | GAATttCCAGGTT TGCCTtGAGG | chr11:108431448 | Not mutated |
| | | GAATGAgCAGGT gTcCCTAGAGG | chr13:56448647 | Not mutated |
| | | GAATGAaCAGtTT TGCCcAGAGG | chr18:53771368 | Not verified |
| | | GAATGcCgAGGcT TGCCTAGCGG | chr18:77550198 | Not mutated |

| | | | | |
|---|---|---|---|---|
| | | GAATGtaCAGaTT TGCCTAGTGG | chr19:246 29132 | Not verified |

**Generation of Eomes and Tbxt null reporter mESC lines**

*Eomes*$^{-/-}$*-GFP* mESC line was generated by transducing *Eomes*$^{-/-}$ mESCs with GFP encoding lentivirus described above. For transduction, ~3 × $10^6$ *Eomes*$^{-/-}$ mESCs were plated on 0.1% gelatin-coated 4-well dishes. After 24h, the growth medium was replaced with fresh medium containing the desired viral dilution and 8µg/mL polybrene. Following a 24h incubation, viral media was removed, and cells were rinsed with PBS and cultured in mESC growth medium. Once confluent, cells were dissociated and reseeded at clonal density. Individual GFP-positive colonies were screened by epifluorescence microscopy (Olympus IX83), manually picked, expanded, and validated for pluripotency and gastruloid-forming potential prior to use in experiments.

Lentiviral tdTomato reporter particles were produced using pUbc tdTomato (Addgene #183710) plasmid using the protocol described above for GFP lentivirus. *Tbxt*$^{-/-}$*-tdTomato* mESC line was generated using a similar method as used for the *Eomes*$^{-/-}$*-GFP* mESC cell line.

All plasmids used in this study are listed in table S2.

**Methods table S2**

| Plasmids | Catalogue number | Source |
|---|---|---|
| pGL3 RARE-d2EGFP | #183055 | Addgene |
| psPAX2 | #12260 | Addgene |
| pMD2.G | #12259 | Addgene |
| pX459 | #62988 | Addgene |
| pUbc tdTomato | #183710 | Addgene |
| pRRL-SIN-PGK-EGFP | #12252 | Addgene |

**Genes used for annotation of clusters in single-cell sequencing data**

**Wild type**

Prechordal plate (*Shh, Gsc, Cer1, Sox2, Sox17, Foxa2, Otx2, Hesx1, Otx2, Dkk1, Six3, Gata4*) ;

Cardiopharyngeal mesoderm (*Dkk1, Mesp1, Fgf10, Eomes, Gsc, Tbx5, Tbx1, Isl1, Hand1, Otx2, Tcf21, Lhx1, Cer1, Fgf5, Pitx2, Sfrp1, Tnnt1, Myl7, Wnt3, Tdgf1*) ;

Pharyngeal mesoderm 1 and 2(*Tcf21, Lhx1, Mesp1, Eomes, Lhx1, Pitx2, Isl1, Tbx1, Gata4, Gata6, Fgf10, Fgf5, Cer1, Otx2*) ;

Anterior paraxial mesoderm (*Isl1, Tbx1, Pitx2, Fgf10, Gata4, Gata6, Sfrp1, Irx5, Tcf15, Meox1, Fst, Foxc2, Uncx, Hoxa2, Hoxb2*) ;

Endoderm *(Sox17, Foxa2, Dkk1, Eomes, Cldn6, Cdh1, Eya1, Six1, Cdx2, Hoxb8) ;*

Node-notochord *(Otx2, Foxa2, Cldn6, Cdh1, Sox2, Hoxa2, Hoxb2, Nodal, Dand5, Foxj1, Shh, Noto, Chrd, T (Tbxt)* ) ;

Caudal epiblast *(Hes7, Shh, T (Tbxt), Sox2, Wnt3a, Wnt3, Tbx6, Rspo3, Nrarp, Mixl1, Chrd, Cdx2, Hoxb8, Hoxb9, Cdh1, Cldn6, Nkx1-2)* ;

Presomitic mesoderm *(Mesp1, Tbx6, Dll1, Pcdh13, Nrarp, Rspo3, Cdx2, Wnt3a, Ripply2, Tcf15, Meox1, Pax3, Foxc2, Hes7, Fst)* ;

Dermomyotome *(Pax3, Pax7, Foxc2, Hes7, Fst, Fli1, Rspo3, Tbx6, Dll1, Pcdh13, Tcf15, Meox1)*

Sclerotome *(Pax1, Pax9, Rspo3, Tbx6, Dll1, Pcdh13, Tcf15, Meox1, Foxc2, Hes7, Fst)* ;

Somite *(Meox1, Tcf15, Pax3, Fst, Pax1, Meox2, Uncx, Dll1, Rspo3, Hoxa2, Hoxb2, Hoxb8, Hoxb9) ;*

Anterior neurectoderm *(Shh, Gsc, Otx2, Sfrp1, Sox2, Sox3, Hesx1, absence of Cdx2 and Hox gene expression)*

Epiblast *(Otx2, Cldn6, Cdh1, Pou5f1, Sox2, Nanog, Dnmt3b, Utf1, Pim2, Slc7a3, Gsh2, Nodal, Tdgf1) ;*

Endothelium *(Nr2f2, Sdk1, Sox17, Kdr, Fli1, Tal1, Etv2, Fev, Sox7) ;*

***Eomes*** $^{-/-}$

Cardiac mesoderm *(Lhx1, Eomes, Gsc, Gata6, Gata4, Fgf10, Hand1, Tbx5, Isl1, Myl7, Ryr2, Ttn, Tnni1, Pitx2, Tcf21, Mesp1) ;*

Neuromesoderm progenitors *(T, Cdx2, Wnt3a, Sox2, Hoxb8, Hoxc8, Hoxb9, Nkx1-2, Epha5, Tbx6, Rspo3, Nrarp, Hes7) ;*

Node-streak border *(Cdx2, T, Sox17, Foxa2, Cldn6, Cdh1, Shh, Chrd, Rspo3, Sdk1, Dall1, Eya1, Dnmt3b) ;*

Mesoderm-primed epiblast *(T, Sox2, Pou5f1, Nanog, Dnmt3b, Utf1, Slc7a3, Cdh1, Otx2, Tdgf1, Ryr2, Tcf15, Nodal, Cldn6, Wnt3, Fgf5, Lhx1) ;*

***Tbxt*** $^{-/-}$

Paraxial mesoderm *(Hand1, Fgf10, Gata4, Gata6, Hoxb8, Hoxc8, Hoxb9, Pitx2, Tbx1, Isl1)* ;

Spinal cord *(Sox3, Nkx6-1, Pax3, Cdx2, Wnt3, Wnt3a, Nkx1-2, Epha5, Hoxb8, Hoxc8, Hoxb9, Sox2, Gbx2, Neurog1)*

Dorsal neural tube *(Nkx6-1, Sox3, Pax3, Cdx2, Wnt3a, Rspo3, Nkx1-2, Epha5, Sox2, Dlx2, Dlx5)*

Neural progenitors *(Neurog1, Sox2, Nkx6-1, Sox3, Pax3, Cdx2)*

Neuromesoderm progenitors *(T, Sox2, Gbx2, Pou5f1, Hoxb8, Hoxc8, Hoxc9, Nkx1-2, Epha5, Cdx2, Tbx6, Wnt3a)*

Neurmesoderm progenitors-neurally committed *(Nkx6-1, Sox3, Pax3, Fst, Cdx2, Nkx1-2, Epha5, Hoxb8, Hoxc8, Hoxc9, Sox2, Tfap2a) ;*

Hindbrain *(Egr2/Krox20, Gbx2, Sox2, Phox2b, Hoxa2, Hoxb2, Pax3, Sox3, Nkx6-1, Tfap2a, Dlx2, Dlx5) ;*

Primed epiblast *(Otx2, Cdh1, Cldn6, Utf1, Pim2, Dnmt3b, Slc7a3, Sox2, Pou5f1, Nanog, Nodal)*
